# Theory of non-dilute binding and surface phase separation applied to membrane-binding proteins

**DOI:** 10.1101/2024.12.31.630850

**Authors:** Xueping Zhao, Daxiao Sun, Giacomo Bartolucci, Anthony A. Hyman, Alf Honigmann, Christoph A. Weber

**Affiliations:** Department of Mathematical Sciences, University of Nottingham Ningbo China; Max Planck Institute of Molecular Cell Biology and Genetics, Dresden, Germany; Technische Universität Dresden, Biotechnologisches Zentrum, Center for Molecular and Cellular Bioengineering (CMCB), Dresden, Germany; Department of Physics, Universitat de Barcelona, Barcelona, Spain; Cluster of Excellence Physics of Life, TU Dresden, Dresden, Germany; Center for Systems Biology Dresden, Dresden, Germany; Faculty of Mathematics, Natural Sciences, and Materials Engineering: Institute of Physics, University of Augsburg, Augsburg, Germany; Max Planck Institute for the Physics of Complex Systems, Dresden, Germany

## Abstract

Surface binding and surface phase separation of cytosolic scaffold proteins on lipid membranes are involved in many cellular processes, such as cell signaling, cell adhesion, and cortex regulation. However, the interplay between surface binding and surface phase separation is poorly understood. In this work, we study this interplay by deriving a general thermodynamic model and applying it to in vitro reconstitution experiments of membrane-binding proteins involved in tight junction initiation. Our theory extends the classical surface binding isotherm to account for non-dilute and heterogeneous conditions where components can phase separate. We use our theory to demonstrate how surface phase separation is governed by the interaction strength among membrane-bound scaffold proteins and their binding affinity to the membrane surface. Comparing the theory to reconstitution experiments, we show that tuning the oligomerization state of the adhesion receptors in the membrane controls surface phase transition and patterning of the scaffold protein ZO1. These findings suggest a fundamental role of the interplay between non-dilute surface binding and surface phase separation in forming the tight junction. More broadly, our work highlights non-dilute surface binding and surface phase separation as a common organizational principle for membrane-associated structures in living cells.

## Introduction

Liquid-liquid phase separation (LLPS) has emerged as a fundamental principle governing the spatial organization of cellular components into dynamic, membrane-less compartments termed biomolecular condensates ***Brangwynne et al. (2011); Mahen and Venkitaraman (2012); Gall (2003); Brangwynne et al. (2009); Fritsch et al. (2021); JR and R. (2009); CJ and R. (2012***). Such condensates create distinct microenvironments that influence a wide range of cellular functions, including signaling ***Su et al. (2016); Wang et al. (2021); Xiao et al. (2022***), transcription ***Boija et al. (2018); Lu et al. (2020***), and stress responses ***Riback et al. (2017***).

While LLPS has been extensively studied in the cytoplasm, protein phase separation is also involved in many biological processes happening at membrane surfaces such as signaling ***Li et al. (2012); CASE et al. (2019); Zeng et al. (2019); Li et al. (2022); Zeng et al. (2018); Su et al. (2016); Zheng et al. (2022***), synapse formation ***Zeng et al. (2018***), cell adhesion ***Beutel et al. (2019); Pombo-García et al. (2024); Wang et al. (2021***) and cortex regulation ***CASE et al. (2019); Wiegand and Hyman (2020***). Protein condensation at membranes is often induced by binding of multivalent scaffold proteins present in the cytoplasm to integral membrane proteins or lipids on the surface of the membrane. Membrane binding of scaffold proteins can result in surface condensation far below the concentration required for phase separation in the bulk, providing spatial-temporal control over the condensation process. For instance, tight junctions are essential for maintaining epithelial barrier integrity by forming continuous adhesive belts around cell perimeters. Recent studies have revealed that the condensation of ZO1 scaffold proteins is a critical step in tight junction assembly ***Beutel et al. (2019***). Specifically, ZO1 initially binds to the receptors in the membrane and undergoes surface condensation at sites of cell-cell contact, a process initiated by the oligomerization of adhesion receptors. These findings underscore the interplay between membrane binding and surface phase separation ***Sun et al. (2024***).

How membrane binding and surface phase separation influence each other is not well understood. Classical surface binding theories, such as the Langmuir isotherm ***Langmuir (1918***), assume dilute concentrations where binding sites are independent and non-interacting. These assumptions are not valid for phase separating systems, where surface densities are *non-dilute* and intermolecular interactions among membrane-bound molecules become important. More recently, non-dilute binding and phase separation have been incorporated into theoretical models ***Zhao et al. (2021, 2024); Liese et al. (2025***). However, these models still rely on simplifying assumptions such as uniform binding sites, which is not matching the reality of biological membranes.

Binding of proteins to biological membranes is typically tightly controlled in space and time by either increasing local densities of binding sites such as in the formation of cell adhesion complexes ***Otani et al. (2019); Zhao et al. (2018b***), or by increasing binding affinities for example by phosphorylation ***Li et al. (2012***). To accurately describe such phenomena, theoretical models must incorporate both non-homogeneous and non-dilute surface densities of proteins at the membrane. While a recent thermodynamic model has linked binding heterogeneity to prewetting phenomena ***Rouches et al. (2021***), developing a theory that integrates these heterogeneous, non-dilute conditions and captures the potential for surface-bound phase transitions remains crucial for a comprehensive understanding of membrane-associated assemblies.

In this work, we present a theoretical framework for the interplay between surface binding and surface phase separation that includes heterogeneous and non-dilute membrane surface densities. We show how the classical binding isotherm must be fundamentally modified when molecules bound to membrane receptors interact strongly and undergo surface condensation. We derived a generalized binding isotherm which extends classical adsorption models by accounting for the non-dilute nature of surface-bound complexes and their interactions. Using this framework, we explored the behaviors of non-dilute surface binding under varying bulk and surface compositions, binding affinities, and interaction strengths between membrane bound complexes. To illustrate the applicability of our theory, we focus on in vitro reconstituted membrane systems that model components of tight junctions. We explain how oligomerization states influence membrane organization. More generally, our theory reveals how non-dilute surface binding and membrane-associated phase separation interact to influence membrane organization, shedding light on a fundamental physicochemical mechanism through which cells regulate membrane associated assembly processes.

### Model for protein binding to receptors on a membrane

In the first part of this work, we discuss a theoretical framework that describes binding of soluble scaffold proteins to lipid membranes via interactions with receptor proteins embedded in a membrane. In addition, the scaffold proteins can undergo (surface) phase separation, which results in formation of protein-dense and protein-dilute regions on the membrane (Fig. 1a,b). The theoretical framework builds upon previous works on the interplay of surface-associated phase transition (surface phase separation, prewetting) and homogeneous surface binding ***Zhao et al. (2021, 2024)***. However, previous models do not involve a receptor component, reducing the stable coexistence of domains poor and rich in bound protein to a specific value of the bulk protein concentration where the phase transition occurs. However, this behavior is not consistent with in vitro experiments where phase coexistence exists over a large range of bulk protein concentrations. In this light, we propose an extended but still minimal model in terms of the number of membrane-associated components: free lipid, free receptor, and receptor bound to a protein from the bulk (Fig. 1a).

**Figure 1.**
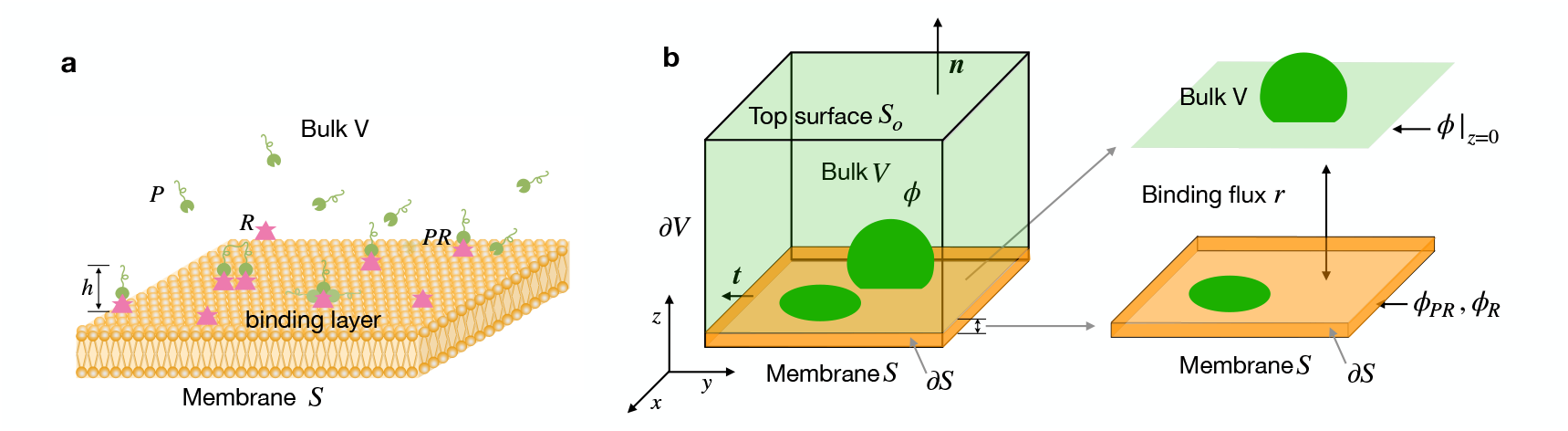
Schematics of the bulk-membrane coupled system. **a** Schematic representation of scaffold proteins binding to the receptors anchored on a lipid bilayer. Each receptor (magenta) is tagged to a lipid (orange) in the supported lipid bilayers (SLBs). Scaffold proteins *P* (green) can bind to receptors *R*, forming a protein-receptor complex *PR*. We introduce two more effective components in the membrane: the lipids with a receptor without any protein bound but a water layer on top, and a lipid patch without a receptor but with the water layer on top. These components are termed “effective” because they also include water molecules that form a molecular layer adjacent to the membrane. For clarity, we do not depict these water molecules in the schematics, thereby providing a clearer view of the surface binding structure. We note that the height of the effective binding layer is denoted as *h*, which is also the height of protein-receptor complex units. Most importantly, the effective components are confined to the membrane and can only move diffusely parallel to it, or unbind and dissolve in the bulk. **b** shows the mathematical notations in our theoretical model. The volume fraction of proteins in the bulk is *ϕ*; *ϕ*_*PR*_, *ϕ*_*R*_ are the area fraction of membrane-bound complexes and free receptors. Moreover, *ϕ*| _*z*=0_ is the volume fraction of protein at the membrane surface, which is denoted as *S. S*_*o*_ indicates the boundary surface of the bulk region *V* opposite to the membrane surface *S*. The boundary surfaces of the bulk region *V*, excluding *S* and *S*_*o*_, are denoted by *∂V* ; The membrane surface’s boundary is represented by *∂S*.

The proteins in bulk can bind to receptors in the membrane, while membrane lipids are not accessible for protein binding. Note that the bulk does not contain any receptors. In our theory, we focus on the classical case of supported lipid bilayers (SLBs); however, we note that the theory can be straightforwardly extended to model real cell membranes even more realistically, e.g. by including more components and the effects of curvature. Since receptors are confined to the membrane, proteins that bind to these receptors can only diffuse within a layer adjacent to the membrane surface. This layer also contains solvent molecules, which are exchanged with the bulk when proteins bind. In the three-dimensional bulk *V*, a binary mixture of protein and solvent molecules is present. To capture processes occurring at the membrane surface, we introduce a quasi-two-dimensional domain *S* representing the membrane environment; see Fig. 1b for an illustration. Moreover, *S*_*o*_ indicates the boundary surface of bulk region *V* opposite to membrane surface *S*. The boundary surfaces of the bulk region *V*, excluding *S* and *S*_*o*_, are denoted by *∂V*. The membrane (one-dimensional) boundary is represented by *∂S*.

To describe the setting outlined above, we propose a spatially coarse-grained model for bulk-membrane coupled systems. To this end, we introduce three effective components on a surface *S*: lipid with receptors bound to protein *PR*, lipid with receptors that are not bound to protein *R*, and lipid with only the water layer on top *L*. Proteins in the bulk are denoted by *P*, and solvent denoted by *W*. For simplicity, we consider an incompressible mixture where the molecular volumes of solvent *v* and protein *v*_*P*_ in bulk and protein-receptor complex 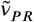 in the membrane are constants. Similarly, the molecular areas of lipid *v*_*L*_, receptor *v*_*R*_ and protein-receptor complex *v*_*PR*_ are constants. The thermodynamics of such an incompressible mixture is characterized by the bulk volume fraction of the proteins *ϕ*(***x***, *t*), and the area fraction of the protein-receptor complex *PR* that is confined in the membrane *ϕ*_*PR*_(***x***_∥_, *t*), and the area fraction, *ϕ*_*R*_(***x***_∥_, *t*), of free receptor *R* on the membrane. Hence, under the incompressibility assumption, the solvent in the bulk occupies the fraction (1 − *ϕ*), and the free lipid in the membrane occupies the fraction (1 − *ϕ*_*R*_ − *ϕ*_*PR*_). All volume fraction fields, in general, depend on position ***x*** = (*x, y, z*) and time *t*. The membrane is located at *z* = 0. We denote *ϕ*| _*z*=0_ as the volume fraction of protein at the boundary surface *S* = {(*x, y, z* = 0) ∈ *V* }. In contrast, the area fraction fields *ϕ*_*PR*_(***x***_∥_, *t*) and *ϕ*_*R*_(***x***_∥_, *t*) depend only on the in-plane coordinates ***x***_∥_ = (*x, y*).

We consider the binding reaction of *α* proteins (labelled by *P*) binding from the bulk to *β* receptors tagged to lipids in the membrane (labelled by *R*); see Fig. 1a for an illustration. This binding process leads to the complex *PR* and the release of (*αn*) solvent molecules (labelled by *W*) to the bulk to conserve the volume of the molecules. The corresponding binding scheme is given as:

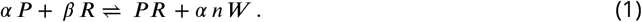

Here, *α* and *β* are the stoichiometric coefficients of the binding process. Volume conservation of the protein implies that the ratio of the molecular volumes between the protein and solvent is *n* = *v*_*P*_ /*v*_*W*_. Note that the receptor *R* is an effective component made of the receptor tagged to lipids and the layer of solvent molecules on top. The height *h* of this layer is set by the complex 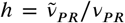, where 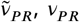 are the molecular volume and area of the protein-receptor complex *PR*, respectively. For volume-conserved binding, the stoichiometric coefficients fulfill 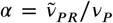 and *β* = *v*_*PR*_/*v*_*R*_.

The binding process is subject to mass conservation that can be expressed in terms of the total particle numbers *N*_*i*_ (*i* = *R, P, PR*):

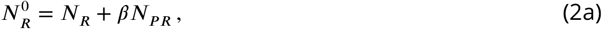

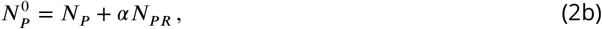

where the superscript 0 indicates the respective total particle numbers conserved in the binding process.

Both conservation laws can be rewritten in terms of the average area fractions 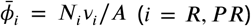 for receptor and complex on the membrane, where *A* is the total area of membrane surface *S*, and the average protein volume fraction in the bulk 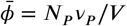:

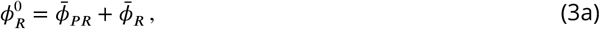

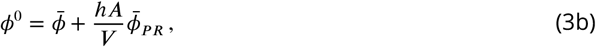

where 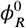 is the conserved total receptor area fraction and 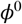 is the conserved total protein volume fraction. The average area fractions are given as 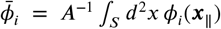 with *i* = *R, P R* and the average protein volume is 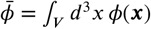.

### Non-equilibrium thermodynamics with non-dilute surface binding

The area fraction fields *ϕ*_*PR*_(***x***_∥_, *t*) and *ϕ*_*R*_(***x***_∥_, *t*) and the bulk protein volume fraction field *ϕ*(***x***, *t*) obey conservation laws for membrane *S* and the bulk *V* :

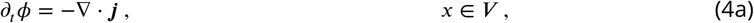

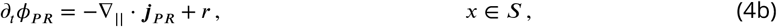

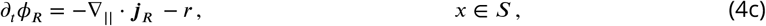

where ***j, j***_*PR*_ and ***j***_*R*_ are the diffusive fluxes of bulk protein *P*, complex *PR* and free receptor *R*. Moreover, *r* denotes the binding rate between bulk and membrane. It is worth noting that *ϕ*_*R*_(***x***_∥_, *t*) denotes the area fraction of free receptors in the membrane. Accounting for the free receptor is crucial for the later comparison to experimental data. Moreover, this additional field is also a key extension compared to previous theoretical model on surface phase separation ***Zhao et al. (2021, 2024)***. The temporal evolution of *ϕ*_*R*_ is governed by Eq. (4c). We define ∇ = (*∂*_*x*_, *∂*_*y*_, *∂*_*z*_) as the gra-dient operator in the three-dimensional bulk and ∇ _||_= (*∂*_*x*_, *∂* _*y*_) on the two-dimensional membrane surface.

Binding relates the protein flux ***n*** · ***j*** normal to the membrane *S* to the binding rate *r*:

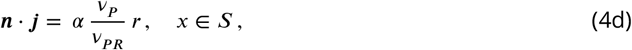

where *α* is the stoichiometric coefficient of the binding protein. The fraction *α v*_*P*_ /*v*_*PR*_ equals the layer height *h*, providing the length scale related to binding from three to two dimensions. The remaining bulk boundaries are considered to be impermeable for all components with the diffusive flux vanishing: ***n*** · ***j*** = 0, *x* ∈ *∂V, S*_*o*_.

The total Helmholtz free energy *F* of the system is composed of different contributions: the free energy in the bulk *V*, the free energy at the membrane surface *S*, and coupling free energy between bulk and membrane. In specific, the total free energy reads:

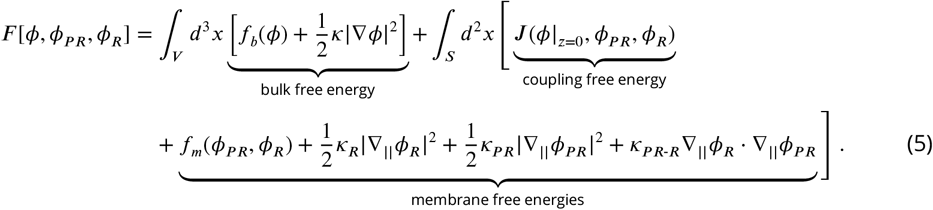

where *f*_*b*_(*ϕ*) is the bulk free energy density in the bulk volume *V, f*_*m*_(*ϕ*_*PR*_, *ϕ*_*R*_) denotes the bulk free energy density at the membrane surface *S*, and *J* (*ϕ* | _*z*=0_, *ϕ*_*PR*_, *ϕ*_*R*_) is the coupling free energy density between bulk and membrane. In addition, we consider the interfacial free energies introduced by the concentration gradients at the phase interfaces, which are characterized by *κ*, and *κ*_*R*_, *κ*_*PR*_, *κ*_*PR*-*R*_, in the bulk and the membrane, respectively.

In the following, we derive the kinetic equations that can relax toward thermodynamic equilibrium. In an isothermal system, the rate of change of the system entropy *S*(*t*) is negatively propositional to the change rate of the total Helmholtz free energy *F* ***Zhao et al. (2024***):

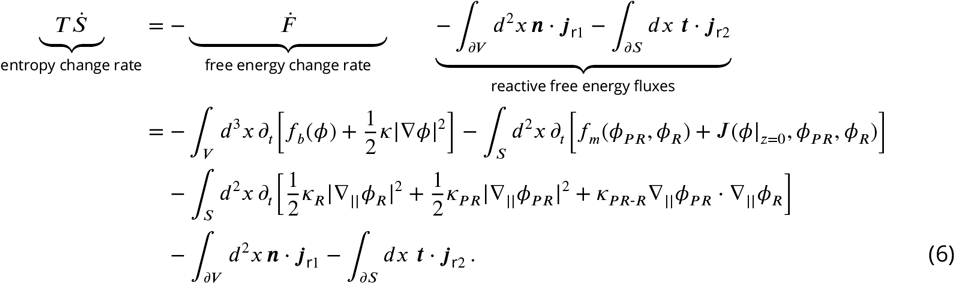

where ***j***_r1_ and ***j***_r2_ denote the non-dissipative, commonly referred to as reactive free energy fluxes. They describe free energy fluxes through the boundaries of bulk and membrane, respectively. Here, ***t*** denotes the normal vector to the boundary *∂S* in the membrane.

Using the conservation laws Eqs. (4), boundary conditions (4d), and defining the reactive free energy fluxes ***j***_r1_ = ***j****μ*/*v*_*P*_ and ***j***_r2_ = ***j***_*PR*_*μ*_*PR*_/*v*_*PR*_ + ***j***_*R*_*μ*_*R*_/*v*_*R*_, we obtain the entropy production rate 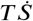 as a product of a pair of thermodynamic fluxes and forces ***Zhao et al. (2024***) as follows:

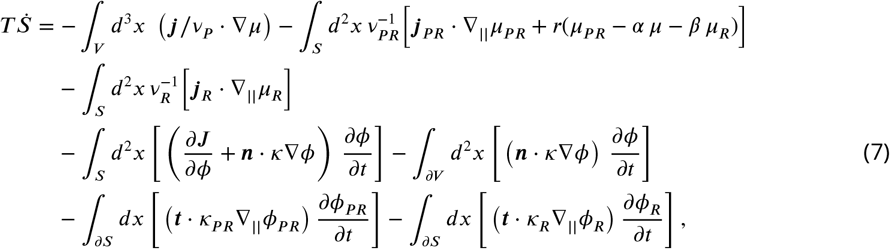

Then, we identify the conjugate thermodynamic fluxes and forces as products in the brackets above.

Consistent with Onsager’s principle and the second law of thermodynamics, we define the constitutive relation between these thermodynamic fluxes and forces to linear order as follows:

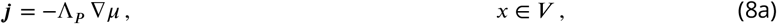

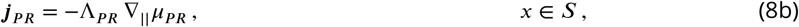

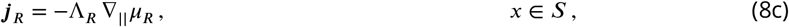

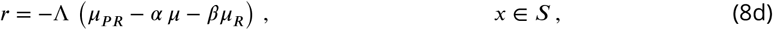

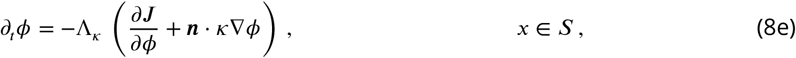

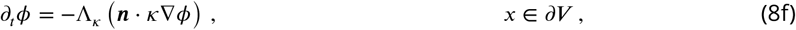

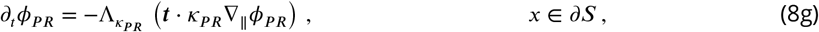

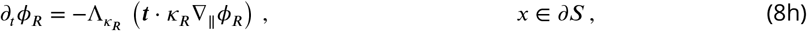

ensuring that the entropy of the system will increase when the system approaches thermodynamic equilibrium. Here, Λ denotes the mobility of binding and Λ_*P*_, Λ_*PR*_ and Λ_*R*_ are the diffusive mobility coefficients in the bulk and the membrane, respectively. Moreover, 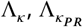 and 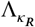 are Onsager coefficients.

Substituting the well-defined fluxes into the conservation laws (4)(a,b,c), we obtain the governing equations of the system as follows:

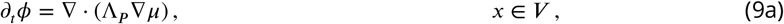

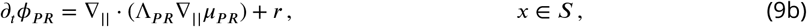

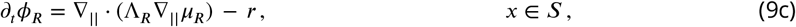

with boundary conditions:

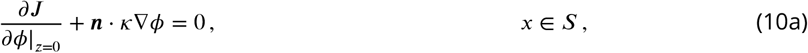

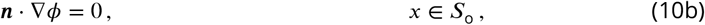

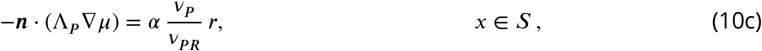

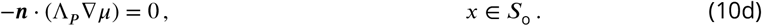

Here, we assume that there are no dissipative fluxes on the boundary *S* when the system approaches the equilibrium state. i.e. we set 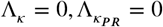 and 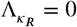. We note that it is possible to have dissipative fluxes ***Zhao et al. (2024***), e.g., −(*∂J*/*∂ϕ*| _*z*=0_ + ***n*** · *κ*∇*ϕ*)Λ_*k*_ = *∂*_*t*_*ϕ* | _*z*=0_, where Λ_*k*_ is the corresponding kinetic coefficient. Here, we focus on the effect of the binding rate *r* leading to compositional changes on a time-scale *τ*, while assuming that other fluxes relax rapidly and remain effectively at equilibrium during the entire kinetic process. Specifically, when Λ_*κ*_*τ* ≫ 1, *∂J* /*∂ϕ*|_*z*=0_ + ***n*** · *κ*∇*ϕ* ≃ 0. To our knowledge, Λ_*κ*_ has not been determined for biological systems.

However, the relaxation is expected to be fast compared to binding processes since the normal concentration gradients ***n*** · ∇*ϕ* are typically confined to microscopic length scales on which diffusion enables quick relaxation.

The boundary condition *∂J* /*∂ϕ*| _*z*=0_+***n***·*κ*∇*ϕ* = 0 in Eq. (10a) characterizes the interactions with the membrane surface and thereby also determine wetting behavior of bulk condensates. The second condition specifies that the opposite side of the membrane *S*_o_ are neutral, i.e., the free energy is not affected by increasing or decreasing the bulk volume fractions adjacent to *S*_o_. Moreover, we did not consider the cross-coupling interaction in the membrane.

We consider the standard mobility scalings,

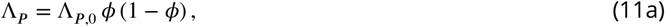

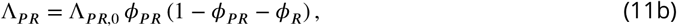

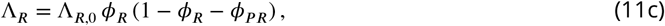

and consider for simplicity the mobility prefactors Λ_*P*,0_, Λ_*PR*,0_, and Λ_*R*,0_ as constants.

The binding rate *r* obeys detailed balance of the rates ***Weber et al. (2019***). However, by Onsager linear response thermodynamics ***Zhao et al. (2024***), we can only obtain the binding rate to linear order in the thermodynamic force Δ*μ* = *μ*_*PR*_ −*β μ*_*R*_ −*αμ*. To account for a general non-linear binding rate that also obeys detailed balance of the rates, we follow Ref. ***Van Kampen (1973***) and write the binding rate

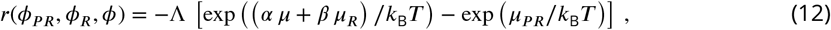

where Λ is the binding rate coefficient. At thermodynamic equilibrium, there are no net flows of matter or energy within the system; that is, the binding flux satisfies *r* = 0.

### Thermodynamic equilibrium with non-dilute surface binding

At thermodynamic equilibrium, the total Helmholtz free energy functional *F* (*ϕ, ϕ*_*P*_, *ϕ*_*R*_, *ϕ*_*PR*_) is minimal. Mathematically, this corresponds to *δF* = 0. In other words, the free energy does not changes in response to small changes of the fields *ϕ, ϕ*_*P*_, *ϕ*_*R*_, *ϕ*_*PR*_ and thus the linear order perturbations *δϕ, δϕ*_*P*_, *δϕ*_*R*_, *δϕ*_*PR*_, of the free energy vanish to zero. The system satisfies two conservation laws: the total number of proteins 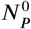 and the total number of receptors at the membrane 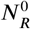 are conserved (Eqs. (2)). Overall, at thermodynamic equilibrium, the first variation of the total Helmholtz free energy *F*, subject to these conservation constraints, vanishes:

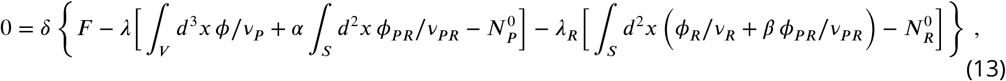

where *λ* and *λ*_*R*_ denote Lagrange multipliers corresponding to the two respective conservation laws. Using Eq. (5), the variation of the total Helmholtz free energy is given as

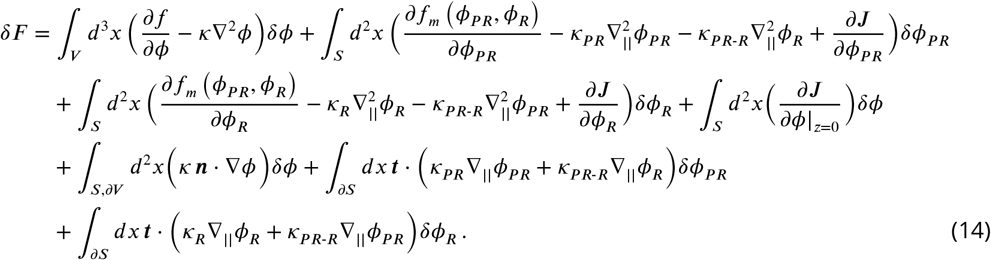

We thus obtain the exchange chemical potentials of the proteins in the bulk *μ*, the protein-receptor complex *μ*_*PR*_ and the free receptor *μ*_*R*_ in the membrane:

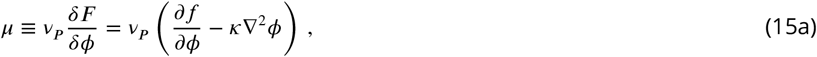

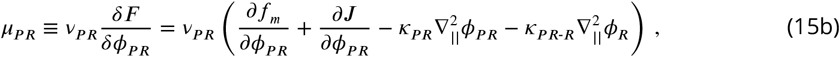

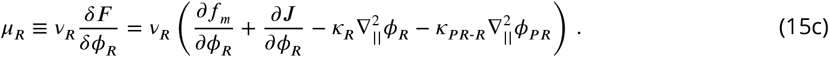

Using Eqs. (15), we find that *μ*_*PR*_ = *α λ*+*β λ*_*R*_, with *λ* = *μ* and *λ*_*R*_ = *μ*_*R*_. As a result, binding equilibrium of the system composed of membrane and bulk is given as:

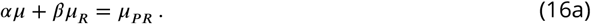

Here, the stoichiometric coefficients *α* and *β* reflect that effective component *PR* is composed of *α* proteins and *β* receptors.

Once the protein binds to the membrane-bound receptor and its local concentration exceeds a certain threshold 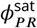, surface phase separation occurs, driven by the attractive interaction between protein–receptor complexes. When two phases, denoted as I and II, coexist in the membrane, phase equilibrium holds:

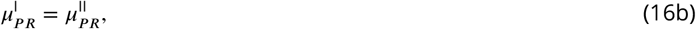

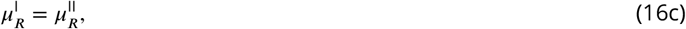

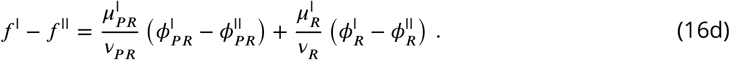

We note that the threshold value for surface phase separation identified above coincides with the volume fraction of the Protein–Receptor complex, i.e., 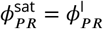. The total amount of receptors in the membrane is conserved (Eq. (3a)) and the conservation for two coexisting phases reads:

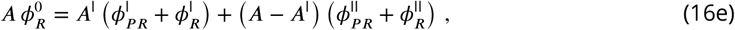

where *A*^I^ is the surface condensate area in the membrane, *A* = *L*^2^ is the total area of the membrane, and 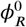 is the initial area fraction of the receptor in the membrane.

We use the boundary conditions at the membrane *S*, the opposite boundary surface *S*_*o*_, and the rest of the boundaries enclosing the bulk, *∂V*, at thermodynamic equilibrium in Eq. (10). Boundary conditions (10) and equilibrium conditions (16) are composite of non-dilute binding equilibrium conditions. In the following section, we will discuss the non-dilute surface binding in detail based on these equilibrium conditions.

### Interplay between surface binding and surface phase separation

#### Non-dilute surface binding

In this section, we derive the general binding isotherm for equilibrium surface binding under non-dilute conditions. Unlike the classical Langmuir model ***Langmuir (1918***), which assumes idealized dilute conditions and no intermolecular interactions, here we incorporate activity coefficients to account for deviations from ideality, such as interactions between different kinds of molecules. This approach leads to a generalized form of the binding isotherm that reduces to the classical Langmuir model under appropriate limiting assumptions.

To capture non-dilute behavior, we express the chemical potentials in terms of standard chemical potentials and activity coefficients:

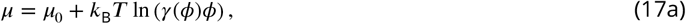

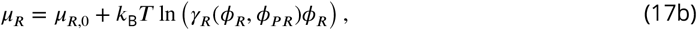

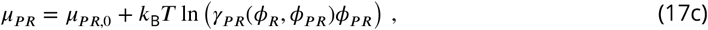

where the terms *γ*(*ϕ*), *γ*_*R*_(*ϕ*_*R*_, *ϕ*_*PR*_), and *γ*_*PR*_(*ϕ*_*R*_, *ϕ*_*PR*_) are activity coefficients capturing non-ideal effects, such as intermolecular interactions or internal energies. The constants *μ*_0_, *μ*_*R*,0_, and *μ*_*PR*,0_ are the corresponding standard chemical potentials that include the internal free energies of the components.

Substituting (17) into the binding equilibrium condition (16a), we rearrange to find,

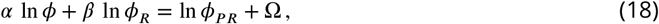

or equivalently,

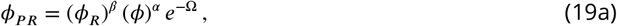

where

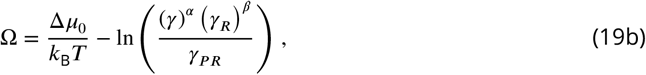

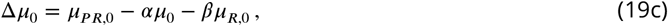

Equation (19a) represents a general non-dilute binding relation, where the effects of interactions at non-dilute conditions are accounted for by activity coefficients *γ, γ*_*R*_ and *γ*_*PR*_. Moreover, Δ*μ*_0_ denotes the differences of standard chemical potential.

To determine the surface coverage, we must also consider the conservation of the total amount of receptor (Eq. (3a)).

By combining this conservation law with the general binding relation (19a), we arrive at a *generalized non-dilute binding isotherm*. Upon choosing a specific form of the free energy densities *f*_*b*_, *f*_*m*_ and *J* in Eq. (5), Eqs. (19) can only be solved numerically due to the non-linear dependence of the activity coefficients on all the area fractions of the components at non-dilute conditions.

For clarity, the non-dilute binding isotherm can be written in a simpler form for the special case of an unimolecular binding process (*α* = *β* = 1). Defining a quantity commonly studied in adsorption isotherms ***Ayawei et al. (2017***), the binding fraction 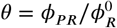, we obtain

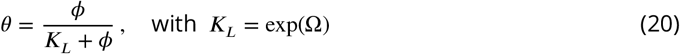

denoting the *generalized Langmuir coefficient*. Here, Ω is the generalized binding affinity given in Eq. (19b), which describes the binding energy per molecule at non-dilute conditions.

In the ideal case where the binding sites are homogeneous and receptors are dilute, the effects of intermolecular interactions in the activity coefficients vanish in Eq. (20). As a result, one obtains the classical Langmuir isotherm ***Langmuir (1918***), and *K*_*L*_ reduces to the Langmuir constant *K*_*L*_ = exp(Δ*μ*_0_/*k*_B_*T*), that is determined solely by the dilute binding affinity Δ*μ*_0_.

In summary, starting from the general equilibrium conditions and incorporating conservation of mass and non-ideal interactions via activity coefficients, we have derived a general binding isotherm applicable to non-dilute conditions (19a). This result not only generalizes the theory of surface binding to non-dilute conditions recovering the classical Langmuir equation in the dilute limit, it also describes phase transitions at the binding surface leading to a heterogeneous distribution of bound molecules.

#### The non-dilute binding isotherm and surface phase separation

In this section, we investigate the intricate behaviors of non-dilute surface binding by varying bulk and surface compositions. Using Flory-Huggins free energy densities for the bulk-membrane system (Appendix 1), we assume unimolecular binding (*α* = *β* = 1) and examine the thermodynamic equilibrium satisfying the conditions of phase coexistence (16b)–(16d) and the generalized binding isotherm (20), we obtain

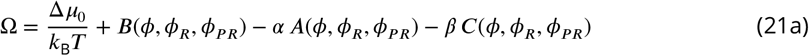

with

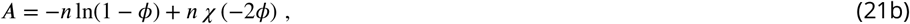

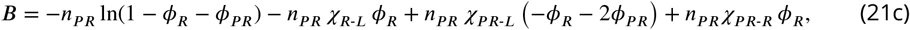

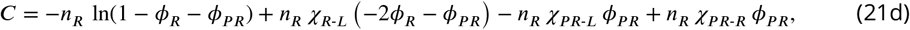

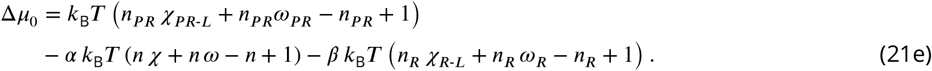

To align with physical constraints and simplify computational implementation, we set the basic parameter values as follows: *ω*_*PR*_ = −1, *χ* = 2.5, *n* = *n*_*PR*_ = *n*_*R*_ = 1, *ω* = *ω*_*R*_ = 0, *χ*_*R*-*L*_ = 0, and *χ*_*PR*-*R*_ = 0. Specifically, *ω*_*PR*_, the internal free energy coefficient, denotes the binding affinity of proteins to the membrane receptors. Setting *ω*_*PR*_ = −1 reflects a moderate attractive interaction, consistent with observed protein-receptor binding in similar systems. The parameter *χ* = 2.5 ensures sufficient phase separation tendencies in the bulk, while *n* = *n*_*PR*_ = *n*_*R*_ = 1 assumes comparable molecular sizes between receptor-bound proteins, unbound receptors, and lipid molecules, simplifying the modeling framework. Finally, since the binary mixture of free receptors and lipids in the membrane does not undergo phase separation, direct receptor–lipid interactions are neglected. Accordingly, we set *χ*_*R*-*L*_ = 0. Due to strong binding in the case of 14mers receptors, free receptors are very dilute in the experiments, making the interaction parameter *χ*_*PR*-*R*_ an irrelevant parameter. Thus, we set *χ*_*PR*-*R*_ = 0 in this study. Finally, setting these interaction parameters to zero allows us to focus on the dominant interactions that determine the system’s binding and phase separation behavior.

We first consider the scenario where phase separation does not occur on the surface, as the attractive interactions between bound proteins (*χ*_*PR*-*L*_ = 2) are insufficient to induce surface phase separation. Under these assumptions, our analysis reveals that the binding fraction *θ* increases monotonically with the bulk protein volume fraction *ϕ* in the non-dilute case (Fig. 2a) This behavior qualitatively resembles the dilute case, which corresponds to a scenario where the total receptor concentration is very low 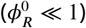; see conservation law (3a). However, in the non-dilute case, the binding fraction *θ* is significantly higher than in the dilute case as the total receptor concentration 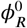 increases. This enhancement is a direct consequence of non-dilute binding. At very high bulk protein volume fractions (*ϕ*), the bulk undergoes phase separation, further altering the system’s behavior.

**Figure 2.**
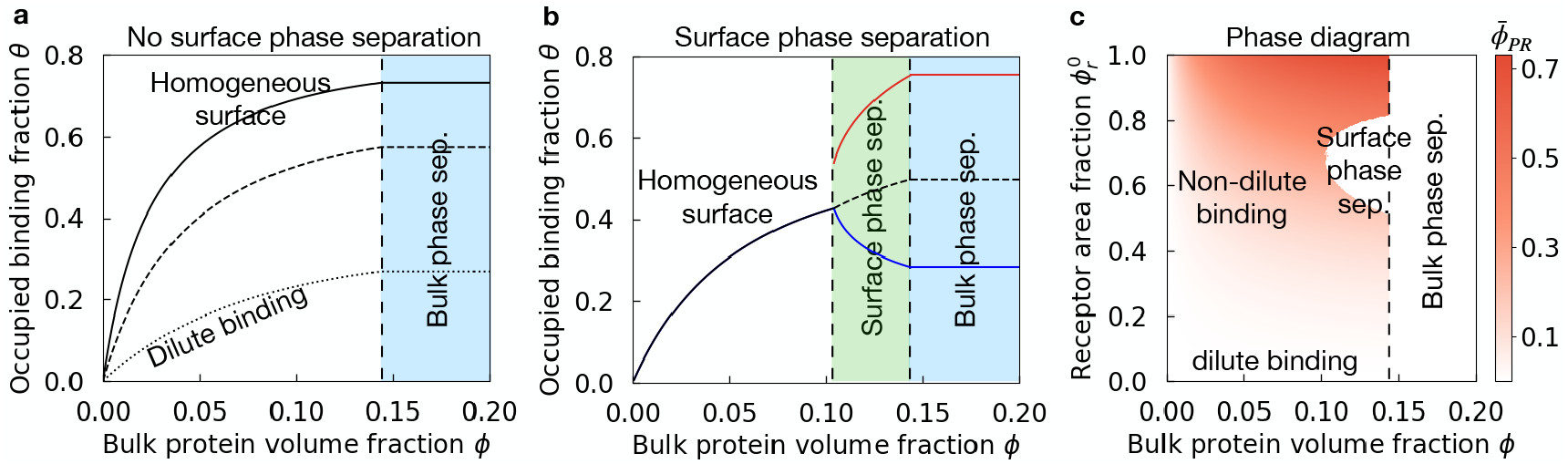
Non-dilute surface binding behaviors with varying surface and bulk compositions. (a) Binding fraction curves 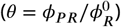 are shown for three different initial receptor area fractions: 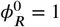 (solid line), 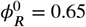 (dashed line) and 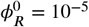 (dotted line), under the assumption of no surface phase separation (*χ*_*PR*-*L*_ = 2). (b) With a stronger interaction strength (*χ*_*PR*-*L*_ = 3.2) and a fixed initial receptor area fraction 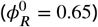, the homogeneous surface mixture (black line) undergoes phase separation into two distinct phases: a protein-rich phase (red line) and a protein-poor phase (blue line) as a function of the intermediate bulk volume fraction, *ϕ*. Additionally, when the bulk protein volume fraction (*ϕ*) exceeds the saturation level, a 3D bulk condensate forms (blue region) that coexist with 2D surface condensates on the membrane. (c) The phase diagram illustrates that intermediate receptor area fractions 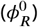 and large bulk protein volume fractions (*ϕ*) promote surface phase separation on the membrane. The basic parameter values are: *ω*_*PR*_ = −1, *χ* = 2.5, *n* = *n*_*PR*_ = *n*_*R*_ = 1, *ω* = *ω*_*R*_ = 0, *χ*_*R*-*L*_ = 0, and *χ*_*PR*-*R*_ = 0.

We now examine the case where the bound proteins at the surface can undergo surface phase separation, forming a protein-rich and a protein-poor phase leading to a heterogeneous distribution of bound proteins *PR* at the surface (see Fig. 2b). A sufficiently large bulk protein concentration is needed to ensure enough proteins are bound to receptors at the surface. Moreover, interactions need to be strong enough (e.g., *χ*_*PR*-*L*_ = 3.2), e.g., the bound proteins attract each other sufficiently strongly.

The effects of both the bulk protein volume fraction *ϕ* and the total amount of receptors on the surface 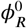 are summarized as a phase diagram in Fig. 2c for a fixed value of *χ*_*PR*-*L*_ = 3.2. It shows that surface phase separation occurs in an intermediate range of both bulk protein and total receptor area fraction 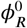. A too-large value of bulk protein volume fraction triggers phase separation in the bulk independent of the receptor and surface binding. Moreover, a too-large amount of receptors on the surface leads to a homogeneous state for volume fraction fractions below bulk phase separation since the mixing entropy on the surface dominates. Similarly, if there are too few receptors on the surface, the enthalpic gains related to the interaction among bound proteins are also outcompeted by the mixing entropy. This overall indicates a reentrant behavior of surface phase separation when varying the total receptor area fraction 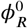.

#### Effects of binding affinity and interactions on surface phase separation

We also investigated how the binding affinity (*ω*_*PR*_) and the interaction strength between membrane-bound proteins (*χ*_*PR*-*L*_) affect surface phase separation. We distinguish two scenarios (Fig. 3a): The three-dimensional bulk is homogeneous, or the bulk is phase-separated.

**Figure 3.**
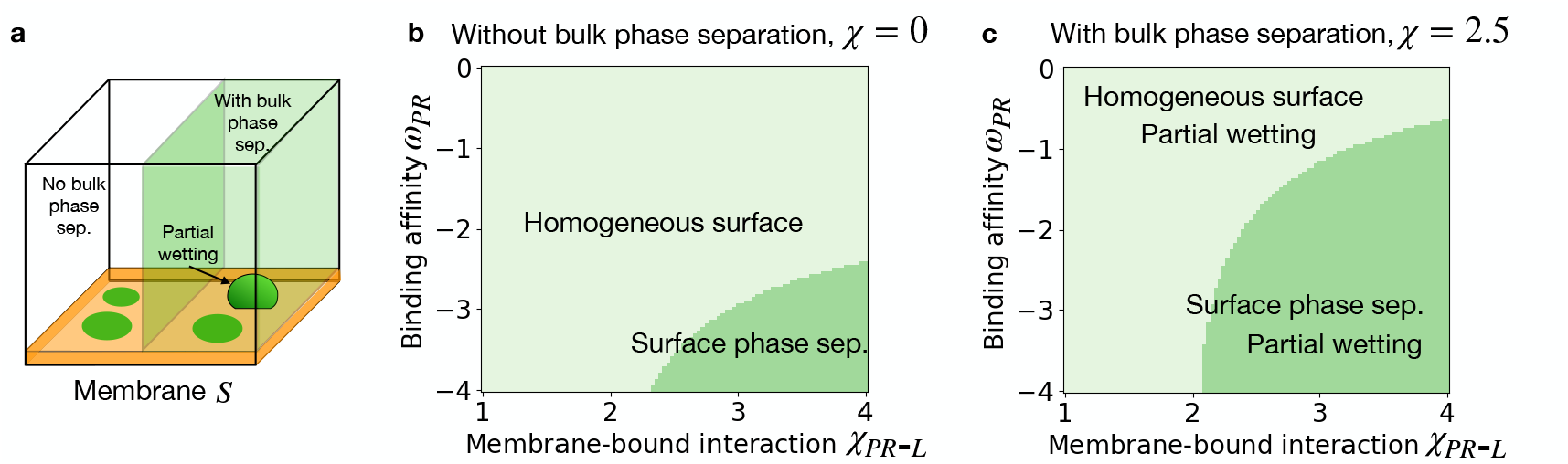
Surface behavior with varying binding affinity and membrane-bound interaction strength. (a) Schematic illustrating bulk-membrane coupled system with surface condensates formed in the membrane *S* without and with bulk phase separation, respectively. To compare phase behavior with and without bulk demixing, we suppress bulk phase separation by setting the protein–solvent interaction to *χ* = 0 and, for each binding affinity *ω*_*PR*_ and membrane–complex interaction strength, compute the surface-binding curve (Fig. 2) to test for surface phase separation. We show the result in phase diagram (b). We then contrast this with a case of stronger bulk interactions (*χ* = 2.5) in (c). If protein interaction in the bulk is strong and thus bulk phase separation occurs. With bulk condensates partially/completely wet on top of the membrane, the surface phase separation occurs in a much larger parameter regime. This illustrates that bulk protein properties, like interaction strength, also regulate the surface phase behavior. The basic parameter values we set in this study are: 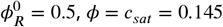, (b) *χ* = 0, (c) *χ* = 2.5. Here, *c*_sat_ represents the bulk volume fraction at the saturation level.

In the absence of bulk phase separation, as shown in Fig. 3b, we find that upon increasing the binding affinity (*ω*_*PR*_) and membrane-bound interaction strength (*χ*_*PR*-*L*_) can induce surface phase separation on the membrane, shifting the system from a homogeneous state to a phase-separated one on the surface. This result underscores the importance of binding affinity and membrane-bound complex interactions as a driving force for surface phase separation.

In the presence of bulk phase separation, as shown in Fig. 3c, we observe a qualitatively similar behavior: strong binding affinity (more negative values of *ω*_*PR*_) and a larger membrane-bound interaction strength (*χ*_*PR*-*L*_) can induce phase separation on the membrane surface. However, the domain of surface phase separation is much larger than the one without bulk phase separation (compare to Fig. 3b), indicating that the bulk interaction parameter *χ* affects phase separation on the membrane. Mechanistically, a larger (and positive) bulk protein interaction strength *χ* increases the local chemical potential of free proteins *μ* near the membrane. As a result, the binding equilibrium (16a) is changed with an enlarged chemical potential difference between unbound and bound states, making more proteins bind from the bulk to the membrane. This leads to the formation of more membrane-bound complexes, making surface phase separation occur over a much broader range of binding affinities and interaction strengths, as illustrated in Figure 3c. In summary, surface phase separation is collectively regulated by the bulk protein interaction strength (*χ*), the binding affinity (*ω*_*PR*_), and the interaction strength among membrane-bound proteins (*χ*_*PR*-*L*_).

Having established the theoretical framework, we now apply it to extract mechanistic insight into tight junction formation, combining with in-vitro reconstitution experiments.

### Mechanism in tight junction formation

Tight junctions are important membrane-associated structures for epithelial and endothelial barrier formation. How surface binding interplays with the condensation process of ZO1 on the plasma membrane at the cell-cell contact sites is not well understood. In this section, we will understand the mechanism for surface phase separation involved in the formation of tight junctions, combining our developed theoretical model and in-vitro reconstitution experiments.

At the cell-cell interface, tight junction receptors including the Claudin family, the CTX family (JAMs) and the MARVEL family (Occludin, Tricellulin, MARVELD3), form multiple *trans-* and *cis-*interactions to mediate cell adhesion (Fig. 4a). Genetic mutation in these *trans-* and *cis-*interactions has been shown to induce tight junction and epithelial barriers deficiency ***Kostrewa et al. (2001); Zhao et al. (2018b); Heinemann and Schuetz (2019***). In this system, we consider ZO1 as the scaffold proteins in the bulk, Claudin2 C terminal (CLDN2-ICD, referred as 1-mer hereafter), which is one of the essential receptors for tight junction function, as the receptors anchored in the membrane. CDLN2-ICD were tagged with His tag and anchored to the membrane with DGS-NTA (Ni) lipid as a mimic of the membrane-associated receptors via His-Ni interaction (Fig. 4b). To mimic the receptor oligomerization happening at the cell-cell contact site, ySMF domain, which is known to form stable 14-mer ***Collins et al. (2003***), was introduced to the CDLN2-ICD construct to get 14-meric receptors (referred as 14-mer hereafter). Scaffold proteins ZO1 were then added to the bulk. See Appendix 4 for methods and materials used in the *in vitro* experiments.

**Figure 4.**
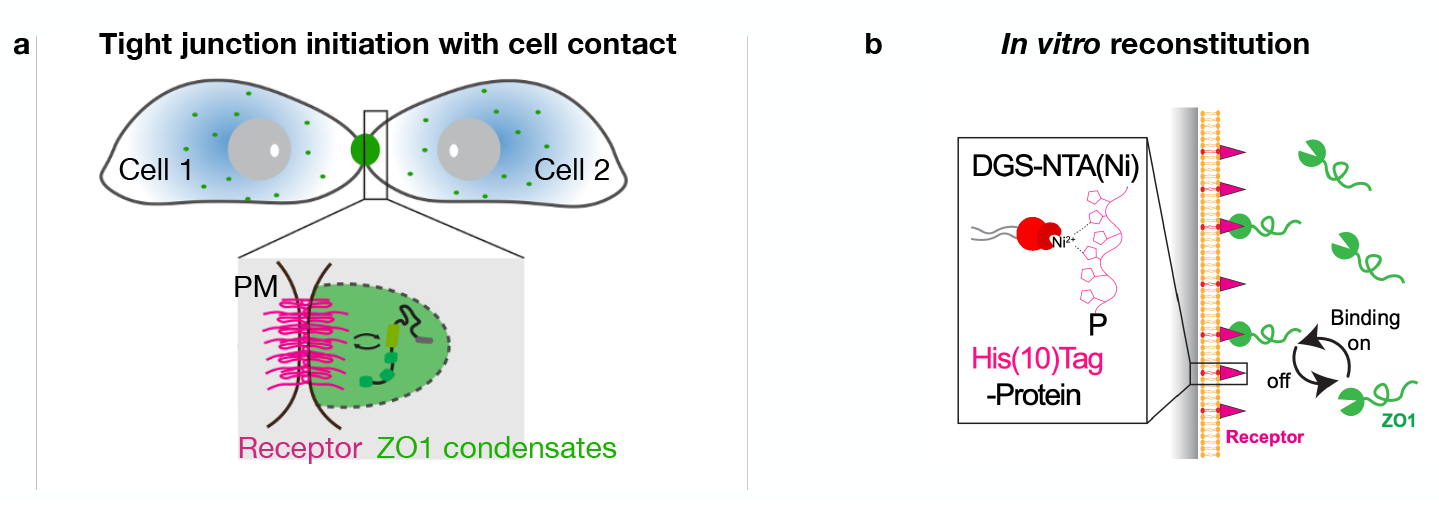
Illustration of tight junction initiation and in vitro reconstitution system. (a) Schematic representation of tight junction initiation during cell-cell contact. Upon contact between Cell 1 and Cell 2, receptors (magenta) on the plasma membrane (PM) interact with scaffold proteins such as ZO1 (green) to form condensates that contribute to tight junction formation ***Beutel et al. (2019***). (b) In vitro reconstitution system showing the binding dynamics of ZO1 proteins (green) with receptors (magenta) on a supported lipid bilayer. The His(10)-tagged receptors are anchored to DGS-NTA (Ni) lipid via His-Ni interaction and can bind and unbind with the ZO1 proteins in the bulk.

Now we use our theory for non-dilute binding to membrane-like surfaces and surface phase separation of bound proteins to model an in vitro reconstitution system for tight junction initiation.

We will show that the theory will be instrumental to characterize binding affinities at non-dilute conditions and elucidate the role of the oligomerization state of the receptor. As a crucial step in this analysis, we estimate the fundamental model parameter values for ZO1 protein binding to claudin receptors by integrating our non-dilute binding theory and experimental data (see Appendix 5 for details). These parameter values, which form the basis of subsequent modeling, are summarized in Table 1.

**Table 1.** Model parameter value and their dimensionless values taken from the experimental data. We note that the molecule sizes *v*_*PR*_, *v*_*R*_ are values for 14-mer receptor. All other parameter values are shared in all the studies.

| Parameter name | Symbol | Value [units] | Value |
| --- | --- | --- | --- |
| **Characteristic scales** |  |  |  |
| Time scale | $t_0 = \Lambda^{-1}$ | 100 [s] | 1 |
| Length scale | $l$ | 10 [$\mu m$] | 1 |
| **Receptor valency** |  |  |  |
| Number of receptors per ZO1 bound | $\beta$ | $\nu_{PR}/\nu_R$ | 1 |
| **Molecular area/volume** |  |  |  |
| Molecular area of free lipid in the membrane | $\nu_L$ | $1.53^2 \pi [nm^2]$ | $1.53^2 \pi / l^2$ |
| Molecular area of bound ZO1 in the membrane | $\nu_{PR}$ | $(9.2/2)^2 \pi [nm^2]$ | $(9.2/2)^2 \pi / l^2$ |
| Molecular volume of solvent in the bulk | $\nu$ | $\frac{4\pi}{3} (9.2/2)^3 [nm^3]$ | $\frac{4\pi}{3} (9.2/2)^3 / l^3$ |
| Molecular volume of scaffold in the bulk | $\nu_P$ | $\frac{4\pi}{3} (9.2/2)^3 [nm^3]$ | $\frac{4\pi}{3} (9.2/2)^3 / l^3$ |
| **Interaction parameter** |  |  |  |
| between free receptor<br>and bound ZO1 in the membrane | $\chi_{PR-R}$ | 0 [$k_B T$] | 0 |
| between free receptor and free lipid | $\chi_{R-L}$ | 0 [$k_B T$] | 0 |
| between ZO1 and solvent in the bulk | $\chi$ | 0 [$k_B T$] | 0 |
| **Conformational coefficient** |  |  |  |
| Gradient coefficient of bound ZO1<br>in the membrane | $\kappa_{PR}$ | $2 \times 10^{-5} [\mu m^{-1}]$ | $2 \times 10^{-4}$ |
| Gradient coefficient of free receptor<br>in the membrane | $\kappa_R$ | $2 \times 10^{-5} [\mu m^{-1}]$ | $2 \times 10^{-4}$ |
| Gradient coefficient of scaffolds in the bulk | $\kappa$ | $2 \times 10^{-5} [\mu m^{-1}]$ | $2 \times 10^{-4}$ |
| **Mobility coefficients** |  |  |  |
| Diffusion coefficient of scaffold in the bulk | $\Lambda_{P,0} k_B T$ | 0.01 [$\mu m^2 s^{-1}$] | 0.01 |
| Diffusion coefficient of bound scaffold<br>in the membrane | $\Lambda_{PR,0} k_B T$ | 0.01 [$\mu m^2 s^{-1}$] | 0.01 |
| Diffusion coefficient of free receptor<br>in the membrane | $\Lambda_{R,0} k_B T$ | 0.01 [$\mu m^2 s^{-1}$] | 0.01 |

#### Receptor oligomerization facilitates surface phase transitions

We analyzed how the oligomerization state of the receptor affects the surface phase transition. In this study, we used two kinds of receptor oligomerization states: 1-mer and 14-mer. By fitting our model (16) to the experimental data (see Appendix 6) for both receptor states (Fig. 5c and 5d and Fig. 6c and 6d), we determined the changes in receptor binding affinity *ω*_*PR*_ and self-interaction parameter *χ*_*PR*-*L*_ of membrane-bound ZO1 complexes. A key finding is that the binding affinity of the 14-meric receptor to ZO1 protein (*ω*_*PR*_ = 3.83, Fig. 6) is much stronger compared to the monomeric receptor (*ω*_*PR*_ = −1.09, Fig. 5). Furthermore, we found that the lower binding affinity of the 1-mer is dominantly caused by a reduced binding fraction, i.e., only a small fraction of 1mer receptors can bind to ZO1 (see Appendix 7 for details); a mechanism previously reported in Ref. ***Erlendsson et al. (2019***). Moreover, from the fits, we obtained that the bound ZO1-14-mer complexes strongly attract each other, i.e., they have a positive self-interaction parameter (Fig. 6). While the fitted self-interaction of the ZO1-1-mer (*χ*_*PR*-*L*_ = −0.4) complex was lower compared to the 14-mer(*χ*_*PR*-*L*_ = 1.09), a wide range of self-interaction values provide similar good fits (blue shade in Appendix 6-fig. 1c), indicating that in the low-affinity binding regime of the 1-mer receptor self-interactions at the surface play a minor role (see Appendix 6-fig. 1c and its related discussion).

**Figure 5.**
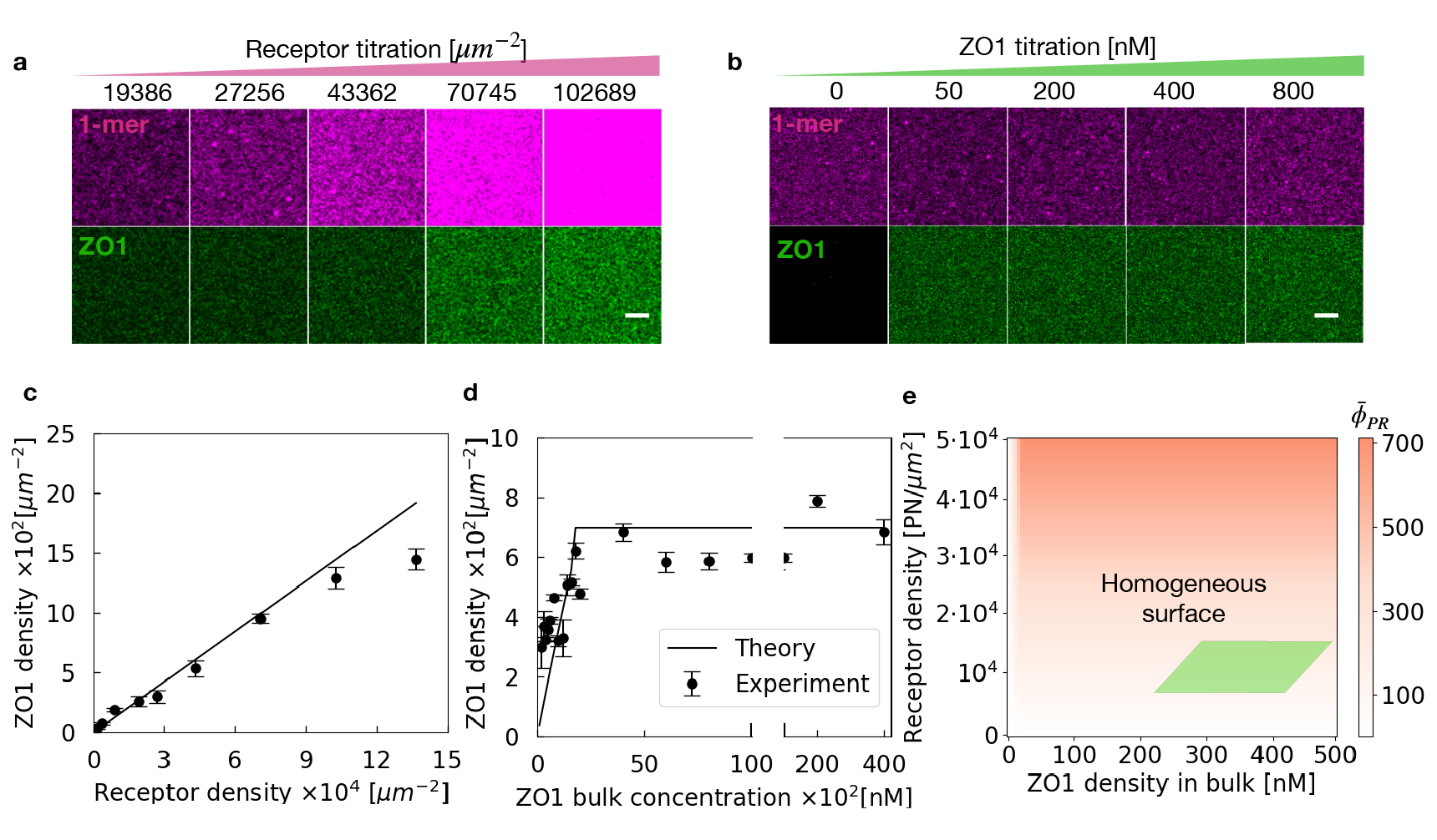
Surface stays homogeneous with 1-mer receptor anchored on the membrane. (a) Fluorescence images of ZO1 and 1-mer at 1-mer receptor titrations, ranging from 19386 to 102689 *μ*m^−2^. (b) Fluorescence images of ZO1 and 1-mer showing the effect of ZO1 titration (0 to 800 nM) on 1-mer binding. Scalar bars in the bottom panels represent 2 *μ*m. (c) The linear relationship between membrane-bound ZO1 density and receptor density on the membrane. This is because, in this study, the binding is saturated for all the receptor densities. In this figure, the dots with error bars represent the experimental data (quantified from figure (a)), and the solid line depicts the theoretical prediction. (d) Curve of membrane-bound ZO1 density as a function of ZO1 bulk density compared between experimental data (points) and theoretical prediction (line). The plateau is obtained from the binding fraction limit for 1-mer, *c* = 0.014. See Appendix 7 for details. The data is a fit to the membrane-bound ZO1 density observed in panel (b). (e) Phase diagram illustrating the homogeneous surface state across a physiological range of receptor and ZO1 densities.

**Figure 6.**
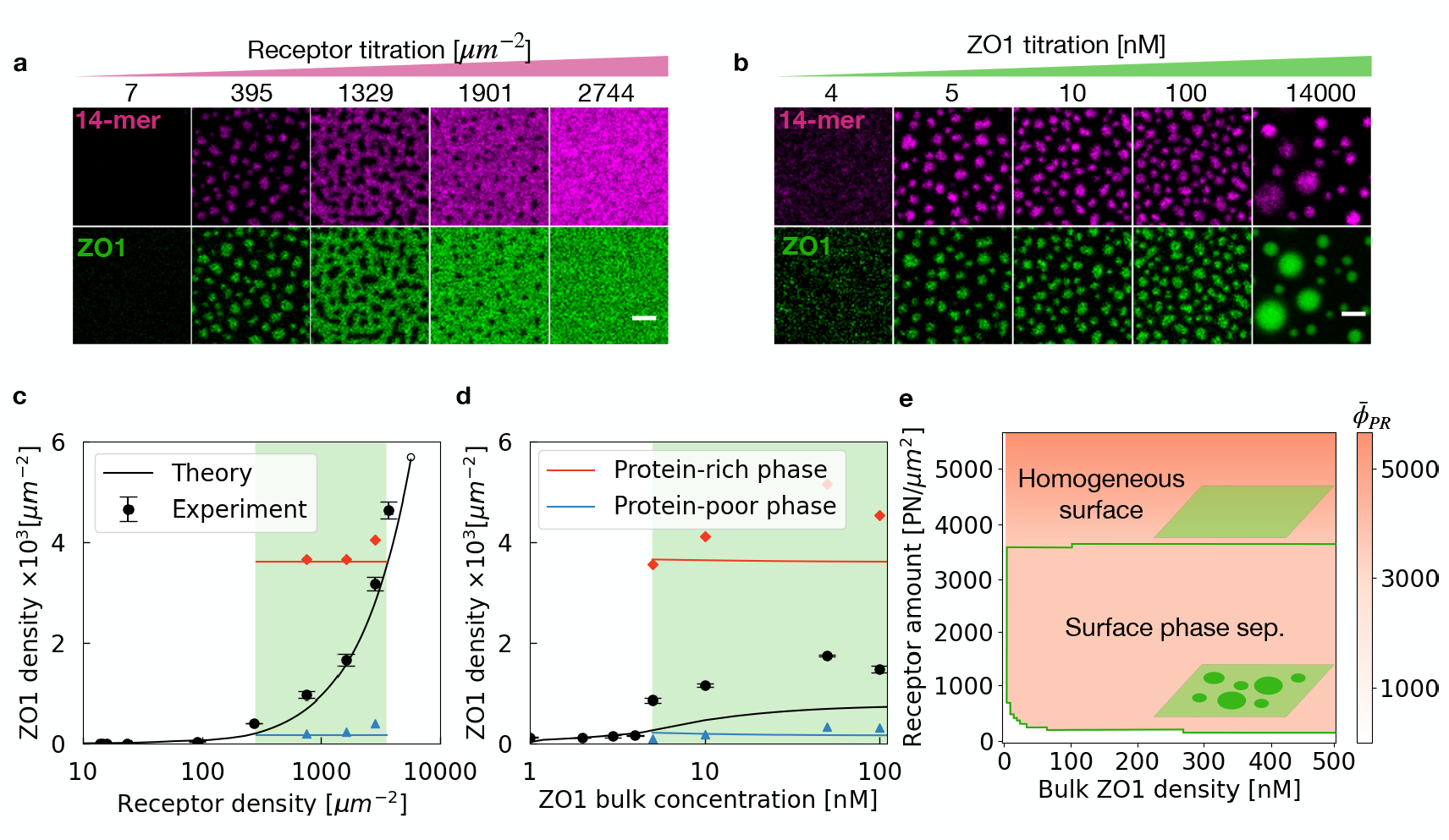
Surface condensates form with 14-mer receptor on the membrane. (a) Fluorescence images showing the distribution of 14-mer receptors and ZO1 across a range of receptor titrations, from 7 to 2744 *μ*m^−2^. Surface phase separation occurs at intermediate receptor densities. Too few or too many receptors result in a homogeneous membrane phase: a dilute phase at low receptor densities and a dense phase at high receptor densities. (b) Fluorescence images of 14-mer receptors and ZO1 as a function of increasing ZO1 bulk concentration, from 4 to 14000 nM, demonstrating the emergence of phase-separated domains. Since receptor density remains constant during this titration, the area of surface condensates does not change with increasing ZO1 bulk concentration. Once the ZO1 bulk concentration exceeds the saturation level, bulk condensates form in the presence of surface condensates (refer to Fig. 3a for illustration). Scalar bars in the bottom panels represent 2 *μ*m. (c) Membrane-bound ZO1 density as a function of receptor density on the membrane, showing how receptor density influences surface phase behavior. In this figure, the dots with error bars represent the experimental data (quantified from figure (a)), and the solid line depicts the theoretical prediction. (d) Membrane-bound ZO1 density dependence on ZO1 bulk concentration, comparing experimental data with theoretical predictions. The data is a fit to the membrane-bound ZO1 density observed in figure (b), revealing a transition from a homogeneous surface phase to a two-phase coexisting state (protein-poor and protein-rich phases). (e) Phase diagram with respect to the receptor density on the membrane and Bulk ZO1 density, indicating that 14-mer receptor promotes surface phase separation.

In summary, our results suggest that receptor oligomerization promotes surface phase separation through two synergistic mechanisms. First, increased binding affinity leads to a higher concentration of ZO1 on the membrane. Second, the strong self-interaction of the protein-receptor complex lowers the saturation concentration required for surface phase separation.

Using the parameters obtained from the fits, we determine the phase diagram for the monomeric (Fig. 5e) and 14-meric receptors (Fig. 6e). These phase diagrams show that there is no surface phase transition for monomeric receptors even for higher ZO1 concentrations, while there is a large region of surface phase separation for the 14-meric receptors.

Based on the agreements between experiments and our thermodynamic model, we conclude that a surface phase transition drives the formation of ZO1-rich surface condensates on membrane-like surfaces. The surface phase transition is controlled by the receptor’s oligomerization state, suggesting cell adhesion-induced oligomerization could act as the key switch for ZO1 surface condensation at cell-cell contacts.

#### Spatial pattern formation of ZO1 surface condensates

When proteins bound to the surface undergo phase separation, multiple condensates form and exhibit a slow ripening kinetics. Theoretical results on surface binding, surface condensates growth and ripening can be obtained from numerically solving the dynamic equations (24) in a three-dimensional bulk volume *V* with surface *S* (Figure 7(a), see Appendix 2 and Appendix 3 for details). Both theoretical predictions and reconstitution experiments reveal an initial stage where surface condensates form and grow rapidly, followed by a prolonged ripening phase characterized by slow kinetics (see Appendix 8, Fig. 1). During this ripening stage, the average condensate areas predicted by the theory align well with experimental observations. Importantly, over a time course of approximately 15 minutes, the amount of bound proteins remains nearly constant as the surface condensates ripen, with only a slight reduction in the number of condensates. To further validate the model, we compared the condensate patterns from experiments and theory at time *t* = 15 min for different receptor titration levels ranging from 25 to 3800 PN/*μm*^2^ and ZO1 titration levels from 4 to 100 nM (Fig. 7(b,c)). The patterns show quantitative agreement between theory and experiments, providing strong evidence that the theoretical framework effectively captures the non-dilute surface binding mechanisms and the condensate formation process on the surface. Furthermore, we observed that the receptor density on the membrane significantly influences the spatial patterns of surface condensates. At lower receptor densities, condensates tend to be sparse and less connected, while higher receptor densities promote larger and more interconnected condensates, consistent with predictions from the theoretical model (Fig. 7(b,c)).

**Figure 7.**
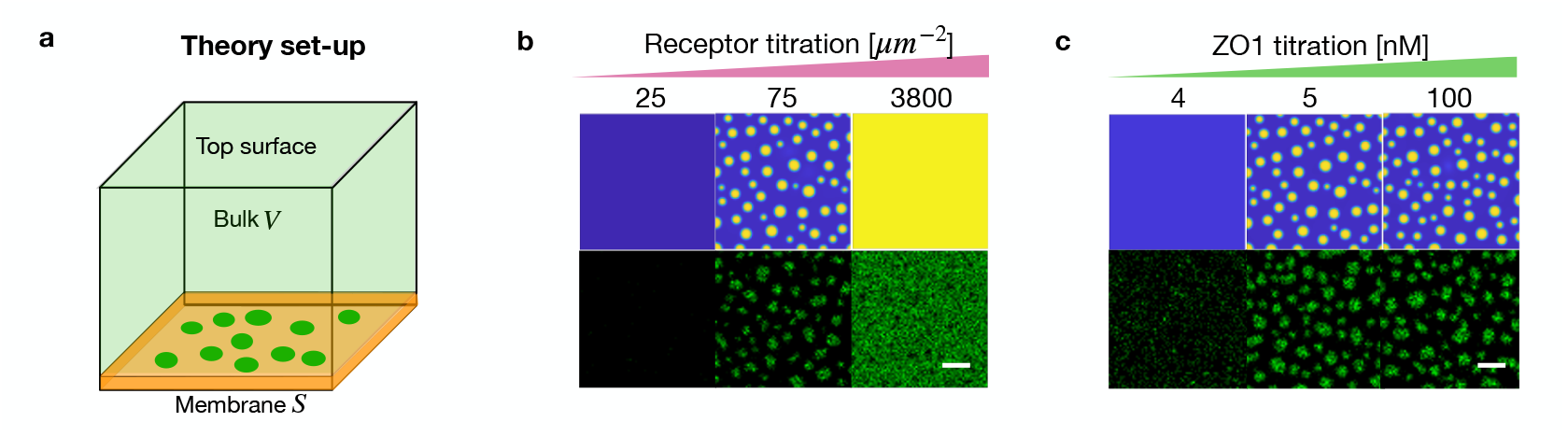
Snapshots of ZO1 surface condensates from simulation (top) and experiments (bottom) when titrating the 14-mer receptor and ZO1 bulk concentration. (a) Schematic of the system in our theoretical model, with 3D bulk volume *V* and membrane surface *S*, where surface condensates formed via surface phase separation. (b) Snapshots of ZO1 from simulation (top row) and experimental observations (bottom) under varying receptor titration levels, from 25 to 3800 *μm*^−2^, at 15 mins. (c) Snapshots of ZO1 from simulation (top row) and experimental observations under varying ZO1 concentration levels, from 4 to 100 nM, at 15 min. The experimental results match well with the theoretical predictions from perspectives of both condensate pattern and condensate size. Scale bars in the bottom panels represent 2 *μm*. we set the diffusivity coefficient value as *D*_*i*_ = 0.01 *μm*^2^*s*^−1^, *i* = *P, P R, R*. The gradient coefficients *κ* = *κ*_*PR*_ = *κ*_*R*_ = 2 × 10^−5^*μm*^−1^.

## Conclusion

In this study, we developed a comprehensive theoretical framework grounded in thermodynamics to describe how soluble scaffold proteins bind lipid membranes through interactions with membrane-embedded receptor proteins. The framework derives a generalized binding isotherm, based on the thermodynamic theory for non-dilute surface binding. This isotherm extends classical adsorption models by accounting for the non-dilute nature of membrane-bound complexes and their interactions. Compared to the classical Langmuir binding isotherm, our theory incorporates interactions among membrane-bound molecules, and accounts explicitly for membrane binding heterogeneity. Consequently, our model is able to capture complex processes such as surface phase separation, which are beyond the descriptive capabilities of classical dilute binding models. Additionally, a kinetic model was derived from non-equilibrium thermodynamics to investigate spatiotemporal patterns of surface condensates.

Using this framework, we explored the behaviors of non-dilute surface binding under varying bulk and surface compositions, binding affinities, and interaction strengths between membrane-bound complexes. The model also enabled us to simulate and analyze surface phase transitions in an in vitro reconstitution system for tight junction initiation. A critical insight from this work is the role of receptor oligomerization in facilitating surface phase transitions, highlighting its importance in membrane-associated condensation phenomena.

Our findings demonstrate that receptor density significantly influences the spatial patterns of surface condensates. At lower receptor densities, condensates are sparse and less connected, whereas higher receptor densities promote larger and more interconnected structures. Moreover, the patterns observed in simulations show quantitative agreement with experimental results, providing strong evidence that the theoretical framework accurately captures the mechanisms of non-dilute surface binding and condensate formation processes, which the dilute model fails to reproduce.

On the membrane with 1-mer receptors, the distribution of membrane-bound proteins remains homogeneous regardless of the bulk ZO1 concentration. The binding isotherm could also be well described by a dilute binding theory, even at non-dilute ZO1 concentrations(see Fig. 5). The reason is that our fits show that the interactions between protein–receptor complexes play only a minor role in this case. In contrast, for 14-mer receptors, surface phase separation emerges at concentrations far below the bulk saturation concentration (see Fig. 6), reflecting both a stronger binding affinity and enhanced intermolecular interactions among protein–receptor complexes on the membrane. Such behavior cannot be captured by the dilute binding isotherm. Therefore, a non-dilute binding theory is essential in our study to account for these complex surface phase behaviors.

In summary, this work bridges theoretical modeling and experimental validation, advancing our understanding of non-dilute surface binding and phase separation on biological membranes. However, it is worth noting that the current theoretical framework has also been applied in limiting cases. For example, in the present model, we looked at a flat membrane, thereby neglecting the effects of membrane curvature, which can significantly influence protein binding and likely also wetting transitions of bimolecular condensates ***Rascón et al. (2018); Lipowsky (2025***). Moreover, we focused on the behavior at and close to equilibrium, and neglected fluxes related to dissipation at system boundaries, such as the triple line. Thus, a richer phenomenology is expected when accounting for far-from-equilibrium dynamics of surface phase separation. Addressing these richer scenarios will be crucial for applying our framework to *in vivo* systems of biomolecular complexes on cellular membrane surfaces.

## Acknowledgments

We thank I. LuValle-Burke for insightful discussions on the binding proteins. We thank Lars Hubatsch on the discussion related to the Tight Juction. We specially thank Frank Jülicher for the collaboration on the fundamental theory on prewetting and surface phase separation (Ref. ***Zhao et al. (2021***, 2024); ***Liese et al. (2025***)). X. Zhao is supported by the FoSE New Researchers Grant, University of Nottingham Ningbo China. D. Sun acknowledges the support from the PEPC facility at MPI-CBG for protein purification. D. Sun was supported by a seed grant of the DFG funded “Physics of Life” Excellent Cluster at TU-Dresden. C. Weber and A. Honigmann acknowledge the SPP 2191 “Molecular Mechanisms of Functional Phase Separation” of the German Science Foundation for financial support and for providing an excellent collaborative environment.

## Appendix 1

### Free energies

To calculate equilibrium conditons (16), we define free energies for the bulk-membrane systems characterizing the energy of the polymer mixtures in the bulk and membrane, respectively.

We use Flory-Huggins free energies in the bulk and membrane, respectively:

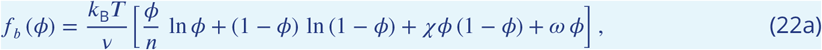

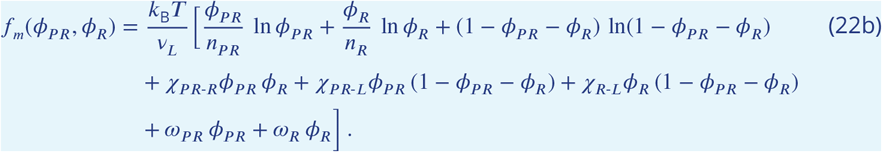

We take the internal free energy of proteins in the bulk as *ω*, denote the internal free energies of protein-receptor complex and free receptor as *ω*_*PR*_ and *ω*_*R*_. We note that *v*_*L*_ represents the area of an effective free lipid patch to avoid considering molecules of very different sizes. As a consequence, to obtain the actual Flory-Huggins parameters corresponding to the lipid-complex interaction, it need to be rescaled by the number of lipids in each lipid patch.

Furthermore, we consider the coupling free energy between the bulk and membrane mixtures. In this study, we focus on the surface phase separation, not the extended prewetting layer or wetting condensates on top of the membrane-bound layer. The coupling free energy only influences on the wetting and prewetting and thus for simplicity, we set coupling free energy as zero in this study. See ***Zhao et al. (2024***) for more complex coupling free energies.

If the free lipid area fraction is zero (*ϕ*_*l*_ = 0), our model reduces to the binary mixture in the model discussed recently ***Zhao et al. (2021***). Without receptors in the membrane (*ϕ*_*R*_ + *ϕ*_*PR*_ = 0), our model yields two decoupled systems: a two-dimensional membrane and a three-dimensional binary mixture in the bulk. If *ϕ*_*l*_ > 0 and *ϕ*_*R*_ > 0, our model corresponds to a ternary mixture in the membrane coupled to a binary mixture in the bulk. This model accounts for one more component, i.e., the protein-receptor complex in comparison to the thermodynamic model developed in the paper ***Rouches et al. (2021***). In addition, our model describes the binding/unbinding events of proteins to the free receptors (or tethers) anchored in the membrane.

We consider a finite system where the bulk has a volume *V* = *L*_*z*_*L*^2^, and *L*_*z*_ denotes the height orthogonal to the membrane surface which has a surface area of *L*^2^. For simplicity, we set *L*_*z*_ = *L* in this study.

## Appendix 2

### Non-dimensionalization of the kinetic model

To solve equations (9) with boundary conditions (10) numerically we rescale length *x* → *x*/*ℓ*, where *ℓ* = 10 *μm* is the experimental field of view, and rescale time *t* → *t* · Λ^−1^. The non-dimensional quantities are:

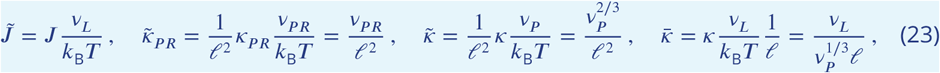

where 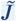 is the non-dimensional binding rate between membrane and bulk. 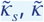, and 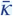 are the dimensional gradient coefficients in the 2D membrane, 3D bulk and bulk boundaries adjacent to the membrane.

We achieve the following non-dimensional equations:

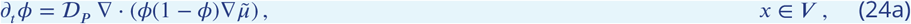

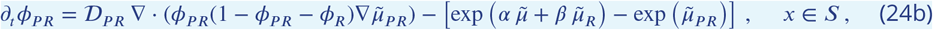

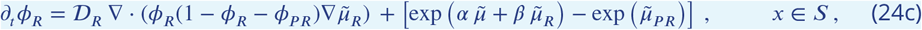

with the non-dimensional boundary equations

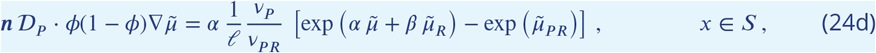

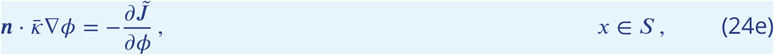

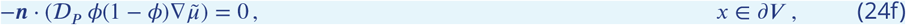

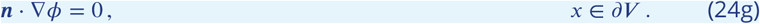

In the equations above, chemical potentials and coupling free energy *J* are measured in units of 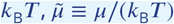 and 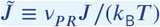. We have also introduced the non-dimensional parameters:

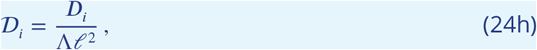

where *i* = *P, P R, R*. Here, 𝒟_*i*_ are inverse “Damköhler numbers”, which compares the time-scale of a chemical reaction, i.e., binding, with the time-scale of diffusive transport. All the parameter values and their dimensionless values in the following study are given in Table 1.

## Appendix 3

### Numerical methods for the kinetic model

In our numerical simulations, the dimensionless governing equations (24) are solved in a three-dimensional domain using a second-order central finite difference method for spatial discretization and the Crank-Nicolson method for time integration. To ensure unconditional energy stability, we apply the Energy Quadratization (EQ) ***Zhao et al. (2018a, 2016); Zhao and Wang (2019***) method, which guarantees that the system’s total energy dissipates while conserving mass throughout the simulation. The computational domain size is set to *L* = 1 in each spatial direction, with *N* = 128 grid points per direction, providing sufficient spatial resolution. The time step is set to Δ*t* = 10^−3^, offering both stability and accuracy during time integration. This setup allows for precise tracking of the system’s kinetics, with a consistent energy dissipation mechanism throughout the evolution of the system.

## Appendix 4

### Methods and materials

#### Protein purification and labeling

##### 1-mer, 14-mer, and ZO1-GFP constructs design

The last C-terminal seven amino acids from human CLDN2 protein was used as the 1-mer receptor for recruiting ZO1 protein to the artificial membranes. To make the 1-mer receptor easier to purify, we fused a SNAP tag on the N terminal to get a recombinant protein SNAP-CLDN2-C7 (referred to as 1-mer). We fused ySMF, a table 14-meric domain, to get a recombinant protein ySMF-CLDN2-C7 (referred to as 14-mer). His tags were used at the N-terminal of both receptors for affinity purification. The human ZO1 sequence was fused with an N-terminal HIS-MBP tag for affinity purification and a C-terminal mGFP tag for visualization.

##### Receptor protein purification

For 1-mer and 14-mer receptors, we used Escherichia coli Rosetta cells to recombinantly express proteins. Proteins were expressed in the Escherichia coli Rosetta cells in LB medium at 16 °C overnight and induced with 0.2 mM IPTG when OD600 reached 0.6 - 0.8. All the following steps were carried out at 4 °C. Cell pellets were collected, washed, and resuspended with a lysis buffer (20 mM Hepes, pH 7.4, 500 mM NaCl, 1 mM MgCl2, 1x protease inhibitor cocktail, 1x benzonase). Cell pellets were broken with LM20 Microfluidizer, 20,000 psi, 2 runs, and clarified by centrifugation at 17,000 g, at 4 °C, for 30 min. Proteins were then purified with metal ion affinity chromatography (Ni-NTA) resin (IMAC, 5mL HiTrap Chelating, GE Healthcare) and followed by size exclusion chromatography with Superdex 200 increase 10/300 GL column on AKTA pure FPLC system (GE). Proteins were collected, aliquoted, and frozen in liquid nitrogen and stored at -80°C in 20mM Hepes, pH 7.4, 500 mM NaCl, 1 mM DTT, and 5% Glycerol.

##### ZO1-GFP protein purification

we used insect cells to recombinantly express ZO1-GFP proteins using the baculovirus expression system. SF9-ESF S. frugiperda cells were infected with ZO1-GFP expression baculoviruses, and then cultured at 27 °C in ESF 921 insect cell culture medium supplemented with 2% fetal bovine serum for 3 days. Cell pellets were then collected, washed, and resuspended with a lysis buffer (20 mM Hepes, pH 7.4, 500 mM NaCl, 1 mM MgCl2, 1 x protease inhibitor cocktail, 1x benzonase). Cell pellets were broken with LM20 Microfluidizer, 5,000 psi, 2 runs, and clarified by centrifugation at 17,000 g, at 4 °C, for 30 min. Proteins were then purified with metal ion affinity chromatography (Ni-NTA) resin (IMAC, 5 mL HiTrap Chelating, GE Healthcare), and followed by amylose resin (NEB). Finally, size exclusion chromatography was performed with the Superose 6 column on the AKTA pure FPLC system (GE). Proteins were collected, aliquoted, and frozen in liquid nitrogen and stored at -80 °C in 20 mM Hepes, pH 7.4, 500 mM NaCl, 1 mM DTT, and 5% Glycerol. His-MBP tag was cleaved and removed with amylose resin before being used for respective assays.

##### Receptor labeling with NHS-ester dye

1-mer and 14-mer receptors were labeled with DyLight 650 NHS Ester (Thermo Scientific, Product No.62262) dye. Highly purified proteins were prepared in 0.05 M sodium borate buffer at pH 8.5 (Thermo Scientific BupH Borate Buffer Packs, Product No. 28384). DyLight NHS Ester dyes were dissolved in DMSO. When labeling the protein, the dye and protein were mixed at a molar ratio of 1:1, and incubated at RT for 1hr. Free dye was removed from the protein by desalting columns (Thermo Scientific Zeba Spin Desalting Columns, Product No. 89882) with buffer containing 20 mM Hepes, pH 7.4, 500 mM NaCl, 1 mM DTT, 5% Glycerol. Fluorescence labeling efficiency was measured with Nanodrop 2000 (ThermoFisher). The labeled and unlabeled proteins were mixed to get a final stock of 5% labeled.

#### In vitro reconstitution of tight junction initiation

##### Liposome preparation

Phospholipids including POPC (Avanti, Product No. 850457), certain amounts of DGS-NTA(Ni) (Avanti, Product No. 790404), 0.1% PEG5000PE (Avanti, Product No. 880230) and 0.1% Rhodamine-PE (Avanti, Product No. 810150) were mixed in chloroform in glass bottles, and dried under vacuum for 2 hrs or overnight. 500 uL buffer (20 mM Hepes, pH 7.4, 150 mM NaCl, and 10 mM MgCl2) was added to the dried lipid film to make a final lipid concentration of 1 mM and resuspended by shaking at a speed of 180 rpm, at 37 °C for 1 hr. The lipid mixture was then transferred to the Eppendorf tube and went through freeze-thaw for 15 runs until the lipid mixture became clear. The liposomes were further clarified by centrifugation at 17,000 g for 30 min and stored at -80°C for long-term storage or 4 °C for use within one week.

##### Supported lipid bilayer (SLBs) preparation

Glass-bottomed 96-well plates (Greiner, Product No. 655891) were pre-cleaned with 2% Hellmanex II overnight and 6 M NaOH for 30 min at RT twice. Before adding liposomes, wells were equilibrated with buffers (20 mM Hepes, pH 7.4, 150 mM NaCl, and 10 mM MgCl2) for 5 min, and left a 60 uL buffer in the well. 20 uL 1 mM liposomes were added and incubated for 20 min. 20 uL 5 M NaCl was then added for another 20 min for the liposomes to further collapse on the glass bottom. Excess liposomes were intensively washed away by pipetting in and out buffers (20 mM Hepes, pH 7.4, 150 mM NaCl, and 10 mM MgCl2) 8 to 10 times. The quality of SLBs was checked by FRAP under a confocal microscope.

##### Surface phase separation assay

SLBs with a certain amount of DGS-NTA(Ni) lipids were blocked with 1 mg/ml BSA for 30 min. 500 nM monomeric or 14-meric receptors were added and incubated for 30 min. Excess receptors were washed away by pipetting in and out buffer (20 mM Hepes, pH 7.4, 150 mM NaCl and 10 mM MgCl2) 8 to 10 times. The images of receptors on SLBs before adding ZO1 were taken with a confocal microscope. The dynamics of receptors on SLBs before adding ZO1 were measured by FRAP. Certain amounts of ZO1-GFP were added for 30 min to form ZO1 surface condensates. 3 images for each well were randomly taken under a confocal microscope. Receptor and ZO1 densities were converted from fluorescence intensity to density numbers with calibration curves obtained with FCS measurement.

##### Fluorescence correlation spectroscopy (FCS)

mGFP and 5% DyLight650 labeled 14-meric receptors were anchored to SLBs via His tag and DGS-NTA(Ni) interaction and excited with 488 nm and 640 nm pulsed laser (Olympus 60x NA = 1.2 water objective). Fluorescence fluctuations were recorded with a time resolution of 500 ns for 10 s. Auto-correlation of the photon traces was performed in MATLAB using a multiple tau correlator. The resulting correlation curves were fitted according to the standard 2D diffusion model including one triplet component using MTALAB (Elson, 2011). The mean particle number N was obtained from the fitting. mGFP and ZO1-mGFP were excited with a 488 nm laser in the bulk. The resulting correlation curves were fitted according to the standard 3D diffusion model including one triplet component using MTALAB (Elson, 2011). The comparison of brightness between the two molecules was calculated by the counts per molecule.

##### Protein density calibration on the membrane with FCS

FCS was performed on SLBs functionalized with different amounts of mGFP and 14-mer-DyLight 650. Mean particle numbers were obtained from FCS fitting. Confocal images were taken for the SLBs functionalized with mGFP and 14-mer-DyLight 650 for extracting the mean fluorescence intensity. The mean fluorescence intensity of mGFP was normalized to ZO1-mGFP by the brightness comparison obtained from FCS measurement in the bulk. Calibration curves were obtained by plotting mean particle numbers as a function of fluorescence mean intensity.

##### Microscopy

Confocal imaging and FCS measurement were performed on a commercial confocal STED microscope (Abberior Instruments, Göttingen, Germany) with pulsed laser excitation (490 nm, 560 nm, 640 nm, 40MHz) and 60x water or 100x oil objectives (Olympus).

##### Quantification and statistical analysis

Images were analyzed with FIJI (https://fiji.sc/) or MATLAB (Mathworks). Statistical details for each experiment can be found in the figure legends and the corresponding methods. Data shown with error bars present the standard deviation (SD). All the experiments were performed more than three times and quantified independently.

## Appendix 5

### Estimation of model parameter for ZO proteins binding to claudin receptors

We analyse the microscope images and compute the density of the components in the membrane and bulk. For this purpose, we define a mask to find the dense and dilute phases, respectively. Specifically, we use a Gaussian filter with a two-dimensional Gaussian smoothing kernel with standard deviation of 0.8 to smooth the raw receptor intensity data. Then, we set the area with *I*_receptor_ ≥ 60 for dense phase(red curve in Appendix 5-fig. 1a), while *I*_receptor_ ≤ 10 for dilute phase(white curve in Appendix 5-fig. 1a). The area in between is the interface area (see Appendix 5-fig. 1). Once we define the dense and dilute phases in the images, we can quantify the density in the dense and dilute phases correspondingly.

**Appendix 5—figure 1.**
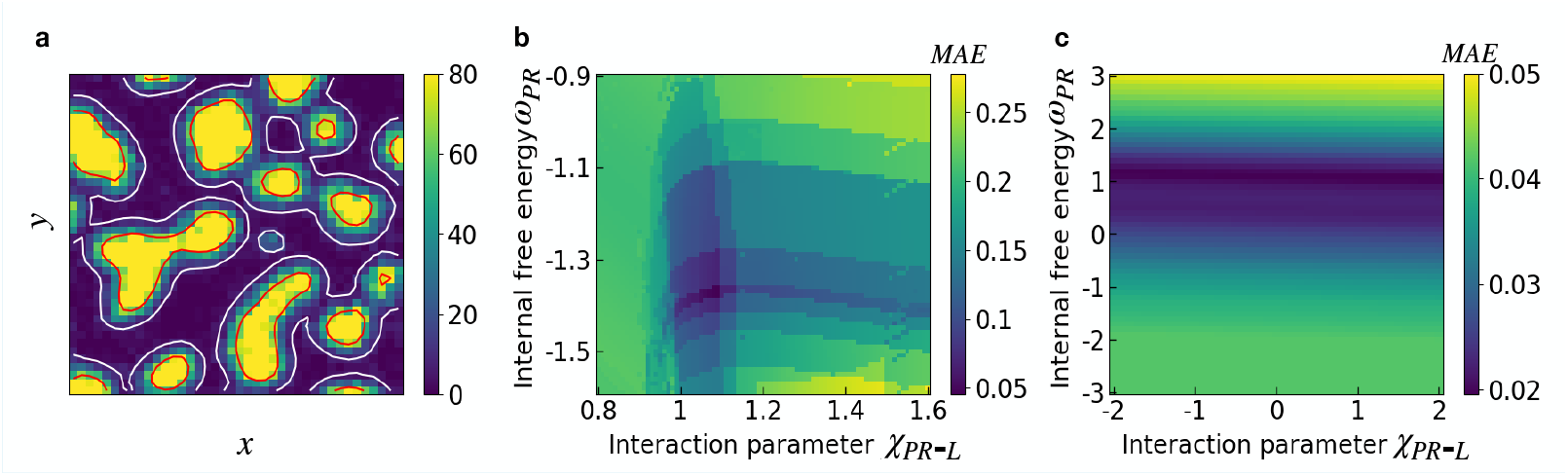
(a) Illustration of the mask for receptor intensity to determine the surface concentrations inside and outside of the surface condensates in the membrane. The blue area represents the dilute phase and the yellow area the dense phase. Mean absolute error (*MAE*) color maps with respect to interaction parameters *χ*_*PR*-*L*_ and internal free energy *ω*_*PR*_ for 1-mer receptor and 14-mer receptor are shown in (b) and (c), respectively. For the considered parameter range, *MAE* is minimized for 14-mer at *χ*_*PR*-*L*_ = 1.09 and *ω*_*PR*_ = −1.37, and for 1-mer at *χ* = −0.4 and *ω*_*PR*_ = 1.1. We note that though there is a minimum of the *MSE* corresponding to at *χ*_*PR*-*L*_ = −0.4, the color map in figure (c) suggests that the *χ*_*PR*-*L*_ value for the 1-mer cannot be determined as there is a “valley” of similar *MAE* value along *χ*_*PR*-*L*_.

According to the experimental studies, membrane phase separation occurs for bulk protein concentrations of a few tens of *nM*. For concentrations in the order of 10 *nM*, there is no phase separation in the bulk since the bulk saturation concentration is about 10 *μM*. Thus, we simplify our model by adapting it in an approximate fashion to these experimental conditions by setting bulk interaction parameter *χ* = 0 and use, for simplicity, equivalent solvent and solute molecule size, i.e., *n* = 1. Furthermore, we choose the coupling free energy equals to zero, implying that the profile along *z, ϕ*(*z*) is constant at thermodynamic equilibrium. Note that during the time evolution, the bulk volume fraction described by Eq. (9a) can be heterogeneous. At thermodynamic equilibrium, however, the bulk can be thought as a finite homogeneous reservoir for proteins.

We roughly estimated the size of 14-mer receptor and 1-mer receptor from the structure in Pymol based on their structures ***Collins et al. (2003); Wilhelm et al. (2021***) as: *d*_14-mer Receptor_ = 9.2 *nm, d*_1-mer Receptor_ = 3.06 *nm*, . The corresponding molecule diameter of the ZO1 protein is achieved from AFM, *d*_ZO1_ = 9.2 *nm*, We choose the size of the lipid patch in the membrane as *d*_lipid_ = 3.06 *nm*. This is an effective free lipid patch to avoid considering molecules of very different sizes as we mentioned before. We note that the molecule area ratios for 14-mer receptor are *n*_*PR*_ = *n*_*R*_ = 9 with lipid patch as a reference component in the membrane, while the ones for 1-mer are *n*_*PR*_ = *n*_*R*_ = 1.

With the given molecule sizes, we determine that the ZO1 bulk volume fraction is exceedingly small (approximately 10^−5^) when the ZO1 number density is 50 *nM* in the bulk. From a numerical standpoint, such a low volume fraction requires a very fine mesh size in both time and space to accurately sample the curvature of the free energy density. This significant curvature originates from the entropic term, and mathematically, from the logarithmic dependence in the free energy. To circumvent the need for extremely fine numerical grids and thus conserve computational resources, we scale the bulk volume fraction *ϕ*^raw^ with a scaling factor such that the scaled volume fraction *ϕ* equals 0.1 when the ZO1 number density is 50 *nM* in the bulk, i.e. *ϕ* = *C*_0_*ϕ*^raw^, where *C*_0_ = 8146.

## Appendix 6

### Model fitting to experimental data of binding ZO1 proteins

Based on the previous discussions, our model has two remaining unknown parameters: the interaction parameter between the protein-receptor complex and free lipid, *χ*_*PR*-*L*_, and the internal free energy difference between the bound and unbound proteins, *ω*_*PR*_. To determine these parameters, we fit our thermodynamic model (Eq. (16)) to experimental data.

We employ two sets of experimental data: one set varies the bulk ZO1 concentration, and the other varies the receptor amount in the membrane. The optimal values of the interaction parameter *χ*_*PR*-*L*_ and the internal free energy coefficient *ω*_*PR*_ are obtained by minimizing the mean absolute error (*MAE*), defined as:

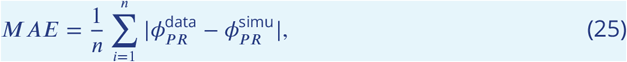

where *n* represents the total number of data points in the experiments. Here, 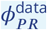 denotes the area fraction of the receptor-ZO1 complex, and 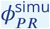 is the simulated area fraction solved from the equilibrium condition (Eq. (16)).

Strikingly, we found very good agreement between the model and the experimental measurements(see Fig. 5c and 5d and Fig. 6c and 6d). In particular, the model recapitulated the experimental ZO1 and receptor (14-mer) concentrations at which surface condensates appear as well as the concentrations in the condensed and dilute phases(see Fig. 6c and 6d). Consistent with the experimental data, the model predicted a jump in the difference in ZO1 concentrations between condensates and their surrounding dilute phase. The binding affinity obtained in the fitting is *ω*_*PR*_ = −1.09 for 1-mer and *ω*_*PR*_ = 3.83 for 14-mer; the interaction strength parameter value is *χ*_*PR*-*L*_ = −0.4 for 1-mer and *χ*_*PR*-*L*_ = 1.09 for 14-mer.

## Appendix 7

### Binding fraction

In the experiments, we observed that maximal fraction of bound receptors is different for different oligomerization states of the receptor, i.e., 1-mer and 14-mer. Specifically, while all 14-mer receptors can be bound by proteins from the bulk, only a fraction of 1-mer receptors are available for binding. This phenomenon was already reported in Ref. ***Erlendsson et al. (2019***) and tracked back to a molecular switch mechanism.

We quantify the binding fraction *c* as the ratio between total membrane-bound protein area fraction and total free receptor area fraction at saturated state, i.e. 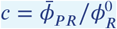. Based on the experimental data, we obtain that *c* = 1 for the 14-mer receptors and *c* = 0.014 for 1-mer receptors. In our model, we include the reduced binding fraction mechanism in the following way:

We calculate the membrane-bound protein area fraction *ϕ*_*PR*_ from the thermodynamic conditions ((16)). Then, (1) If 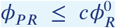, a finite binding fraction *c* has no effects. (2) If 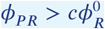, we use

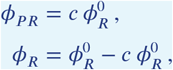

where 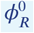 is the average area fraction of the free receptors in the membrane initially.

## Appendix 8

### Kinetic properties of the surface phase separation

**Appendix 8—figure 1.**
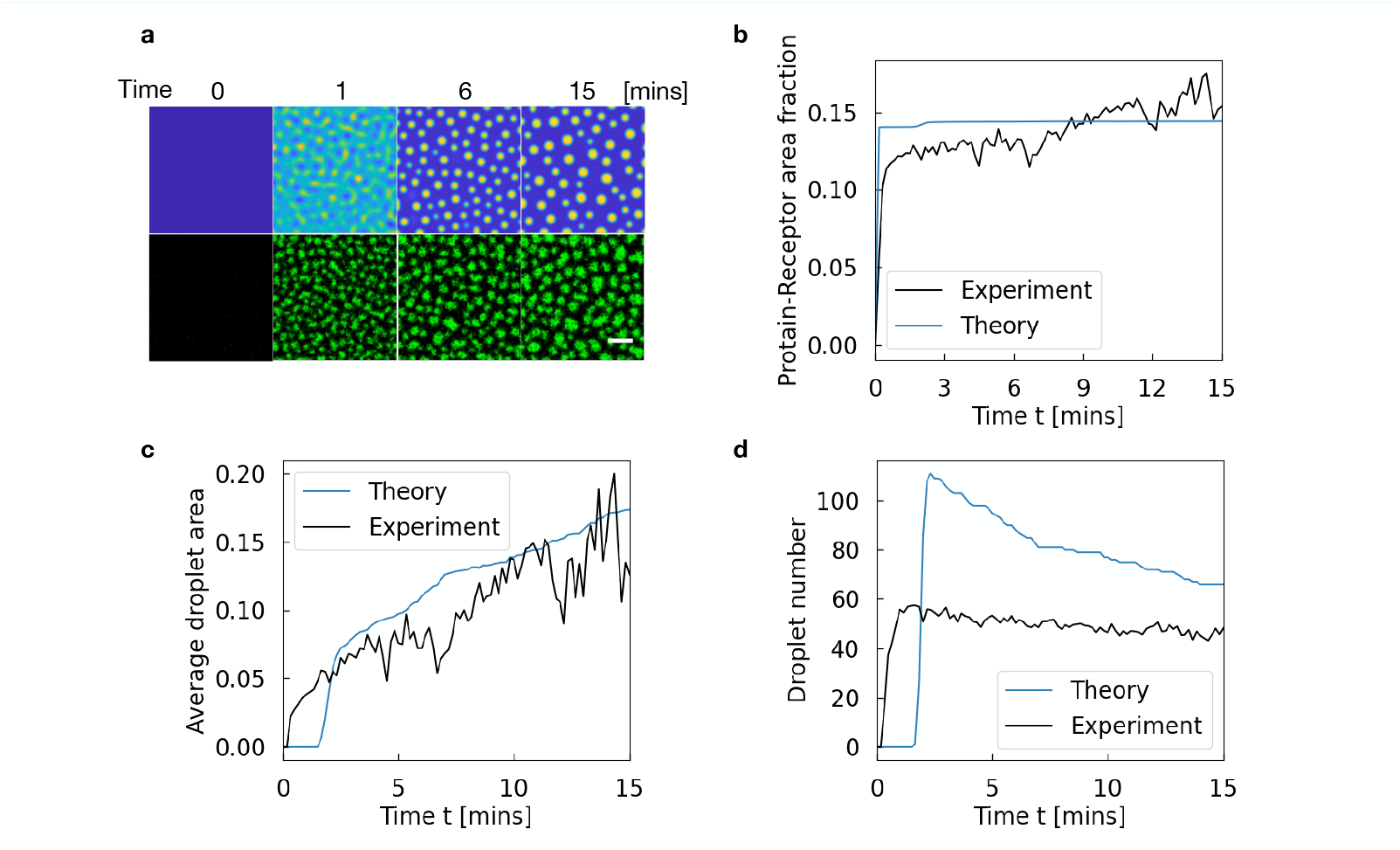
(a) Time-resolved snapshots illustrating the emergence and growth of ZO1-rich condensates on a model membrane surface (top row: theoretical simulation; bottom row: experimental observation). Initially uniform at *t* = 0, the membrane surface progressively develops well-defined droplet patterns by 15 minutes. The system includes 100 nM ZO1 in the bulk and 800 *μm*^−2^ 14-mer Receptor in the membrane. The scale bar in the experimental images represents 2 *μm*. (b) Temporal evolution of the total membrane-bound ZO1 fraction, comparing the theoretical prediction (blue line) and experimental measurement (black line). Both approaches show an increase over time, and they closely agree on the final coverage levels. (c) Average droplet area as a function of time, demonstrating the consistency between theory (blue line) and experiment (black line) in capturing the size growth and stabilization of condensates. (d) Time-dependent count of individual droplets, showing that both the model (blue line) and the experimental data (black line) reproduce the overall trend in droplet formation and coalescence. we set the diffusivity coefficient value as *D*_*i*_ = 0.01 *μm*^2^*s*^−1^, *i* = *P, P R, R*. The gradient coefficients *κ* = *κ*_*PR*_ = *κ*_*R*_ = 2 × 10^−5^*μm*^−1^.

We compare the simulated formation and evolution of ZO1-rich regions on the membrane (top row) to those observed experimentally (bottom row) (see Fig. 1). At the start, both simulation and experiment show a relatively uniform distribution of receptors and ZO1. Over time, small clusters begin to appear and grow more distinct, and by 15 minutes both systems feature well-defined, separated condensates.

The overall pattern of how these clusters develop is quite similar in both simulation and experiment. We observe that surface condensates rapidly form once the critical concentrations of receptor-ZO1 complex are exceeded. However, after this initial rapid phase, the system transitions to much slower coarsening kinetics (Appendix 8-fig. 1, movie S1). This slow-down in coarsening kinetics is consistent with the experimental data and can be explained theoretically by two possible mechanisms. First, the depletion of receptors in the dilute phase due to co-condensation of receptors and scaffold proteins likely reduces the ripening fluxes between surface condensates (Appendix 8-fig. 1a). This reduction in flux slows down the further growth and coalescence of condensates, effectively stalling the kinetic process. Alternatively, the slow-down may be attributed to an ageing process ***Jawerth et al. (2020***) of the surface condensates and could be explained by the limited movement of surface condensates on the membrane with the gain of mass ***Snead et al. (2022***).

These results illustrate that the theoretical framework, incorporating appropriate diffusivity and interaction parameters, closely reproduces not only the qualitative spatial patterns observed experimentally but also key quantitative descriptors of ZO1 droplet dynamics, including membrane coverage(see Fig. 7b, Fig. 7c and Appendix 8-fig. 1a), droplet size(see Appendix 8-fig. 1c), and droplet number(see Appendix 8-fig. 1d). However, the timing of the initial binding and recruitment does not fully align with the experimental observations shown in Appendix 8-fig. 1(b). This discrepancy may also stem from differences between the model’s assumed initial conditions and those in the experimental setup, as well as the limited spatiotemporal resolution of the experimental measurements. These factors lie beyond the scope of this manuscript. Nevertheless, the strong overall agreement indicates that our model provides a valuable representation of the key mechanisms behind ZO1 surface condensate formation.

## References

Ayawei N, Ebelegi AN, Wankasi D. Modelling and interpretation of adsorption isotherms. Journal of Chemistry. 2017; 2017(1):3039817. doi: 10.1155/2017/3039817.

Beutel O, Maraspini R, Pombo-García K, Martin-Lemaitre C, Honigmann A. Phase separation of zonula occludens proteins drives formation of tight junctions. Cell. 2019 10; 179:923–936.e11. doi: 10.1016/j.cell.2019.10.011.

Boija A, Klein IA, Sabari BR, Dall’Agnese A, Coffey EL, Zamudio AV, Li CH, Shrinivas K, Manteiga JC, Hannett NM, Abraham BJ, Afeyan LK, Guo YE, Rimel JK, Fant CB, Schuijers J, Lee TI, Taatjes DJ, Young RA. Transcription factors activate genes through the phase-separation capacity of their activation domains. Cell. 2018 12; 175. doi: 10.1016/j.cell.2018.10.042.

Brangwynne CP, Eckmann CR, Courson DS, Rybarska A, Hoege C, Gharakhani J, Jülicher F, Hyman AA. Germline P granules are liquid droplets that localize by controlled dissolution/condensation. Science. 2009 5; 324(5935):1729–1732. doi: 10.1126/science.1172046.

Brangwynne CP, Mitchison TJ A HA. Active liquid-like behavior of nucleoli determines their size and shape in Xenopus laevis oocytes. Proceedings of the National Academy of Sciences. 2011 1; 108(11):4334–4339. doi: 10.1073/pnas.1017150108.

Case LB, Zhang X, Ditlev JA, Rosen MK. Stoichiometry controls activity of phase-separated clusters of actin signaling proteins. Science. 2019; 363(6431):1093–1097. doi: 10.1126/science.aau6313.

CJ D, R P. P-bodies and stress granules: possible roles in the control of translation and mRNA degradation. Cold Spring Harbor Perspectives in Biology. 2012 7; 4. doi: 10.1101/cshperspect.a012286.

Collins BM, Cubeddu L, Naidoo N, Harrop SJ, Kornfeld GD, Dawes IW, Curmi PMG, Mabbutt BC. Homomeric ring assemblies of eukaryotic Sm proteins have affinity for both RNA and DNA: crystal structure of an oligomeric complex of yeast SmF*. Journal of Biological Chemistry. 2003; 278(19):17291–17298. doi: 10.1074/jbc.M211826200.

Erlendsson S, Thorsen TS, Vauquelin G, Ammendrup-Johnsen I, Wirth V, Martinez KL, Teilum K, Gether U, Madsen KL. Mechanisms of PDZ domain scaffold assembly illuminated by use of supported cell membrane sheets. eLife. 2019 1; 8:e39180. doi: 10.7554/eLife.39180.

Fritsch AW, Diaz-Delgadillo AF, Adame-Arana O, Hoege C, Mittasch M, Kreysing M, Leaver M, Hyman AA, Jülicher F, Weber CA. Local thermodynamics govern formation and dissolution of Caenorhabditis elegans P granule condensates. Proceedings of the National Academy of Sciences. 2021 7; 118(37):e2102772118. doi: 10.1073/pnas.2102772118.

Gall JG. The centennial of the Cajal body. Nature Reviews Molecular Cell Biology. 2003 12; 4:975–980. doi: 10.1038/nrm1262.

Heinemann U, Schuetz A. Structural features of tight-junction proteins. International Journal of Molecular Sciences. 2019; 20(23):6020. doi: 10.3390/ijms20236020.

Jawerth L, Fischer-Friedrich E, Saha S, Wang J, Franzmann T, Zhang X, Sachweh J, Ruer M, Ijavi M, Saha S, Mahamid J, Hyman AA, Jülicher F. Protein condensates as aging Maxwell fluids. Science. 2020; 370(6522):1317–1323. doi: 10.1126/science.aaw4951.

JR B, R P. Eukaryotic stress granules: the ins and outs of translation. Molecular Cell. 2009 12; 36(6):932–941. doi: 10.1016/j.molcel.2009.11.020.

Kostrewa D, Brockhaus M, D’Arcy A, Dale GE, Nelboeck P, Schmid G, Mueller F, Bazzoni G, Dejana E, Bartfai T, Winkler FK, Hennig M. X-ray structure of junctional adhesion molecule: structural basis for homophilic adhesion via a novel dimerization motif. The EMBO Journal. 2001; 20(16):4391–4398. doi: 10.1093/emboj/20.16.4391.

Langmuir I. The adsorption of gases on plane surfaces of glass, mica and platinum. Journal of the American Chemical Society. 1918 6; 40(9):1361–1403. doi: 10.1021/ja02242a004.

Li P, Banjade S, Cheng HC, Kim S, Chen B, Guo L, Llaguno M, Hollingsworth JV, King DS, Banani SF, Russo PS, Jiang QX, Nixon BT, Rosen MK. Phase transitions in the assembly of multivalent signalling proteins. Nature. 2012; 483(7389):336–340. doi: 10.1038/nature10879.

Li R, Li T, Lu G, Cao Z, Chen B, Wang Y, Du J, Li P. Programming cell-surface signaling by phase-separation-controlled compartmentalization. Nature Chemical Biology. 2022 11; 18(12):1351–1360. doi: 10.1038/s41589-022-01192-3.

Liese S, Zhao X, Weber CA, Jülicher F. Chemically active wetting. Proceedings of the National Academy of Sciences of the United States of America. 2025; 122:e2403083122. doi: 10.1073/pnas.2403083122.

Lipowsky R. Complex remodeling of biomembranes and vesicles by condensate droplets. Soft Matter. 2025; 21:7370–7392. doi: 10.1039/D5SM00585J.

Lu Y, Wu T, Gutman O, Lu H, Zhou Q, Henis YI, Luo K. Phase separation of TAZ compartmentalizes the transcription machinery to promote gene expression. Nature Cell Biology. 2020 3; 22(4):453–464. doi: 10.1038/s41556-020-0485-0.

Mahen R, Venkitaraman AR. Pattern formation in centrosome assembly. Current Opinion in Cell Biology. 2012 2; 24(1):14–23. doi: 10.1016/j.ceb.2011.12.012.

Otani T, Nguyen TP, Tokuda S, Sugihara K, Sugawara T, Furuse K, Miura T, Ebnet K, Furuse M. Claudins and JAM-A coordinately regulate tight junction formation and epithelial polarity. Journal of Cell Biology. 2019 08; 218(10):3372–3396. doi: 10.1083/jcb.201812157.

Pombo-García K, Adame-Arana O, Martin-Lemaitre C, Jülicher F, Honigmann A. Membrane prewetting by condensates promotes tight-junction belt formation. Nature. 2024 8; 632:647–655. doi: 10.1038/s41586-024-07726-0.

Rascón C, Pausch J, Parry AO. First-order wedge wetting revisited. Soft Matter. 2018; 14:2835–2845. doi: 10.1039/C8SM00342D.

Riback JA, Katanski CD, Kear-Scott JL, Pilipenko EV, Rojek AE, Sosnick TR, Drummond DA. Stress-triggered phase separation is an adaptive, evolutionarily tuned response. Cell. 2017 3; 168:1028–1040. doi: 10.1016/j.cell.2017.02.027.

Rouches M, Veatch SL, Machta BB. Surface densities prewet a near-critical membrane. Proceedings of the National Academy of Sciences. 2021; 118(40):e2103401118. doi: 10.1073/pnas.2103401118.

Snead WT, Jalihal AP, Gerbich TM, Fritsch AW, Higgs HN, Gladfelter AS. Membrane surfaces regulate assembly of ribonucleoprotein condensates. Nature Cell Biology. 2022; 24:461–470. doi: 10.1038/s41556-022-00882-3.

Su X, Ditlev JA, Hui E, Xing W, Banjade S, Okrut J, King DS, Taunton J, Rosen MK, Vale RD. Phase separation of signaling molecules promotes T cell receptor signal transduction. Science. 2016 4; 352(6285):595–599. doi: 10.1126/science.aad9964.

Sun D, Zhao X, Wiegand T, Martin-Lemaitre C, Borianne T, Kleinschmidt L, Grill SW, Hyman AA, Weber C, Honigmann A. Assembly of tight junction belts by ZO1 surface condensation and local actin polymerization. Developmental Cell. 2024; 60:1234–1250.e6. doi: 10.1016/j.devcel.2024.12.012.

Van Kampen N. Nonlinear irreversible processes. Physica. 1973; 67(1):1–22.

Wang B, Zhang L, Dai T, Qin Z, Huasong Lu LZ, Zhou F. Liquid–liquid phase separation in human health and diseases. Signal Transduction and Targeted Therapy. 2021 8; 6(1):290. doi: 10.1038/s41392-021-00678-1.

Weber CA, Zwicker D, Jülicher F, Lee CF. Physics of active emulsions. Reports on Progress in Physics. 2019 4; 82(6):064601. doi: 10.1088/1361-6633/ab052b.

Wiegand T, Hyman AA. Drops and fibers — how biomolecular condensates and cytoskeletal filaments influence each other. Emerging Topics in Life Sciences. 2020 10; 4(3):247–261. doi: 10.1042/ETLS20190174.

Wilhelm J, Kühn S, Tarnawski M, Gotthard G, Tünnermann J, Tänzer T, Karpenko J, Mertes N, Xue L, Uhrig U, Reinstein J, Hiblot J, Johnsson K. Kinetic and structural characterization of the self-labeling protein tags HaloTag7, SNAP-tag, and CLIP-tag. Biochemistry. 2021 8; 60(33):2560–2575. doi: 10.1021/acs.biochem.1c00258.

Xiao Q, McAtee CK, Su X. Phase separation in immune signalling. Nature Reviews Immunology. 2022 4; 22(3):188–199. doi: 10.1038/s41577-021-00572-5.

Zeng M, Chen X, Guan D, Xu J, Wu H, Tong P, Zhang M. Reconstituted Postsynaptic Density as a Molecular Platform for Understanding Synapse Formation and Plasticity. Cell. 2018; 174:1172–1187.e16. doi: 10.1016/j.cell.2018.06.047.

Zeng M, Díaz-Alonso J, Ye F, Chen X, Xu J, Ji Z, Nicoll RA, Zhang M. Phase separation-mediated TARP/MAGUK complex condensation and AMPA receptor synaptic transmission. Neuron. 2019 11; 104:529–543.e6. doi: 10.1016/j.neuron.2019.08.001.

Zhao J, Yang X, Li J, Wang Q. Energy stable numerical schemes for a hydrodynamic model of Nematic liquid crystals. SIAM Journal on Scientific Computing. 2016; 38(5):A3264–A3290. doi: 10.1137/15M1024093.

Zhao J, Yang X, Gong Y, Zhao X, Yang X, Li J, Wang Q. A general strategy for numerical approximationss of non-equilibrium models-part I: thermodynamic systems. International Journal of Numerical Analysis & Modeling. 2018 8; 15(6):884–918. http://global-sci.org/intro/article_detail/ijnam/12613.html.

Zhao J, Krystofiak ES, Ballesteros A, Cui R, Van Itallie CM, Anderson JM, Fenollar-Ferrer C, Kachar B. Multiple claudin–claudin cis interfaces are required for tight junction strand formation and inherent flexibility. Communications Biology. 2018; 1. doi: 10.1038/s42003-018-0051-5.

Zhao X, Wang Q. A second order fully-discrete linear energy stable scheme for a binary compressible viscous fluid model. Journal of Computational Physics. 2019 10; 395:382–409. doi: 10.1016/j.jcp.2019.06.030.

Zhao X, Bartolucci G, Honigmann A, Jülicher F, Weber CA. Thermodynamics of wetting, prewetting and surface phase transitions with surface binding. New Journal of Physics. 2021 12; 23:123003. doi: 10.1088/1367-2630/ac320b.

Zhao X, Liese S, Honigmann A, Jülicher F, Weber CA. Theory of wetting dynamics with surface binding. New Journal of Physics. 2024 9; 26:103025. doi: 10.1088/1367-2630/ad80bb.

Zheng Q, Chen Y, Chen D, Zhao H, Feng Y, Meng Q, Zhao Y, Zhang H. Calcium transients on the ER surface trigger liquid-liquid phase separation of FIP200 to specify autophagosome initiation sites. Cell. 2022 10; 185(22):4082–4098.e22. doi: 10.1016/j.cell.2022.09.001.

